# Nuclear envelope reshaping around the vacuole determines the morphology of the ribosomal DNA

**DOI:** 10.1101/2021.06.23.449658

**Authors:** Emiliano Matos-Perdomo, Silvia Santana-Sosa, Jessel Ayra-Plasencia, Félix Machín

## Abstract

The ribosomal DNA array (rDNA) of *Saccharomyces cerevisiae* has served as a model to address chromosome organization. In cells arrested before anaphase (mid-M), the rDNA acquires a highly structured chromosomal organization referred to as the rDNA loop, whose length can double the cell diameter. Previous works established that complexes such as condensin and cohesin are essential to attain this structure. Here, we report that the rDNA loop adopts distinct morphologies that arise as spatial adaptations to changes in the nuclear morphology triggered during the mid-M arrests. Interestingly, the formation of the rDNA loop results in the appearance of a space under the loop (SUL) which is devoid of any nuclear component yet colocalizes with the vacuole. We finally show that the formation and maintenance of the rDNA loop and the SUL require TORC1 and membrane synthesis. We propose that the rDNA-associated nuclear envelope (NE) reshapes into a loop to accommodate the vacuole, with the nucleus becoming bilobed.

## Introduction

One of the most remarkable visual events in cell biology is chromosome condensation, which takes place when cells transit from G2 into prophase and metaphase (M phase). In higher eukaryotes, chromosomes are condensed over a central scaffold formed by topoisomerase II alpha, condensin I and condensin II ^1^. Condensation requires the reorganization of the chromatin fiber into nested loops ^1–3^, so that chromosomes progressively shorten their length while becoming wider. The formation and expansion of these loops continues until chromosomes acquire their characteristic rod-like appearance by late metaphase.

The yeast *Saccharomyces cerevisiae* has served as an instrumental model for understanding fundamental processes of the eukaryotic cell. Its powerful genetic tools enable precise characterization of protein function. Condensin (there is a unique condensin complex in yeast) has not been an exception, and has been scrutinized extensively ^4–8^. However, whereas in higher eukaryotes chromosome condensation is cytologically evident, this is not in yeast, where most of the nuclear mass remains visually amorphous throughout the cell cycle. The only exception to this is the ribosomal DNA array (rDNA), located on the right arm of chromosome XII (cXIIr), and whose repetitive nature facilitates its visualization by fluorescence in situ hybridization (FISH) and fluorescence microscopy through specific rDNA binding proteins (e.g., Net1, Fob1, Cdc14, etc.) tagged with GFP variants. Previous studies showed that the rDNA of wild type cells appears unstructured in interphase, with a spotted and diffuse morphology by FISH, referred to as puff, and a crescent shape at the nuclear periphery when labelled with GFP-tagged coating proteins ^8–12^. As cells enter G2/M, this disorganized rDNA becomes a highly organized bar-like structure. In cells arrested in late metaphase (sometimes referred to as mid-M in yeast) by the microtubule-depolymerizing drug nocodazole (Nz), the rDNA bar bends out of the rest of the chromosomal mass to become a horseshoe-like loop. This rDNA loop has been considered a condensed state of the repetitive locus and depends on active condensin for its establishment and maintenance ^5,8,13^.

Condensin is not the only factor involved in the establishment and maintenance of the rDNA loop. Cohesin, which keeps sister chromatids together until anaphase, and the Polo-like kinase Cdc5 are also involved, yet their role is thought to regulate condensin activity on the rDNA ^14–16^. Besides these, we previously reported that the rDNA loop requires an active Target of Rapamycin Complex 1 (TORC1) ^12^. TORC1 is the master complex that regulates cell growth and metabolism, controlling anabolic processes in the cell such as ribosome biogenesis, protein synthesis and lipid synthesis ^17–19^. Previous works, including ours, have shown that TORC1 inactivation impinges on the morphology of the rDNA/nucleolus, reducing its size both in interphase and mitosis ^12,20,21^. Several mechanisms under the control of TORC1 have been proposed for this, including inhibition of both ribosome biogenesis and rDNA transcription, and autophagy of nucleolar components into the vacuole ^22–24^. Because transcription of rDNA genes demands high levels of energy and resources, it is not surprising that the rDNA physiology is a main target of TORC1 ^25,26^.

Yeasts undergo a closed mitosis and do not disassemble the nuclear envelope (NE) when entering M-phase ^27^. The rDNA is tethered to the NE throughout the cell cycle ^28,29^. Importantly, previous works showed that both a prolonged mid-M arrest and mutants related to lipid metabolism led to an enlargement of the NE that specifically affected the membrane region associated with the nucleolus and where the rDNA attaches to ^30–33^. Here, we show that the vacuole, which occupies a large proportion of the cell volume, serves as a template to reconfigure the nuclear morphology during NE expansion in mid-M, and thus emerges as a major determinant of the morphology of the rDNA loop. The NE acquires tubular projections that contain the rDNA and distal parts of chromosome XII. These projections often bend themselves around the vacuole and reshape the nucleus towards a bilobed morphology, with one lobe containing most of the nuclear mass, the other one just the distal regions of chromosome XII, and with the nucleoplasmic handle that connects both lobes forming the rDNA loop. We further show that this reorganization of the nuclear shape requires an active TORC1 and new membrane synthesis, but is independent on reported nuclear-vacuole contacts. We discuss how our new findings affect our vision of the rDNA loop as a model of chromosome condensation.

## Results

By FISH, the mid-M (Nz-arrested) rDNA loop appears as a handle that bends out of the rest of chromosomes and makes the chromosomal mass to resemble a handbag ^8,9,11^. The same pattern is often seen under the microscope when the rDNA is labelled with Net1-GFP and the nuclear mass with DAPI (Figure 1A) ^10,12^. Unlike FISH, fluorescence microscopy enables to see the loop in the context of an entire cell. Two features stand out above all and called our attention. Firstly, the loop goes across a significant proportion of the cell space; and secondly, it leaves a space under the loop (SUL) that is as large as the rest of the nuclear mass stained with DAPI, at the very least. Thus, both the loop and the SUL often occupy a remarkable cell area, far beyond the proportional expectations for the nuclear/cytoplasmic volume ratios ^34,35^. At first, this suggests that either this volume ratio is dramatically shifted towards the nucleus or the nucleus is flattened in order to occupy a larger area and thus accommodate the enlarged rDNA loop. Hence, we aimed to study both the loop and the SUL in detail.

**Figure 1.**
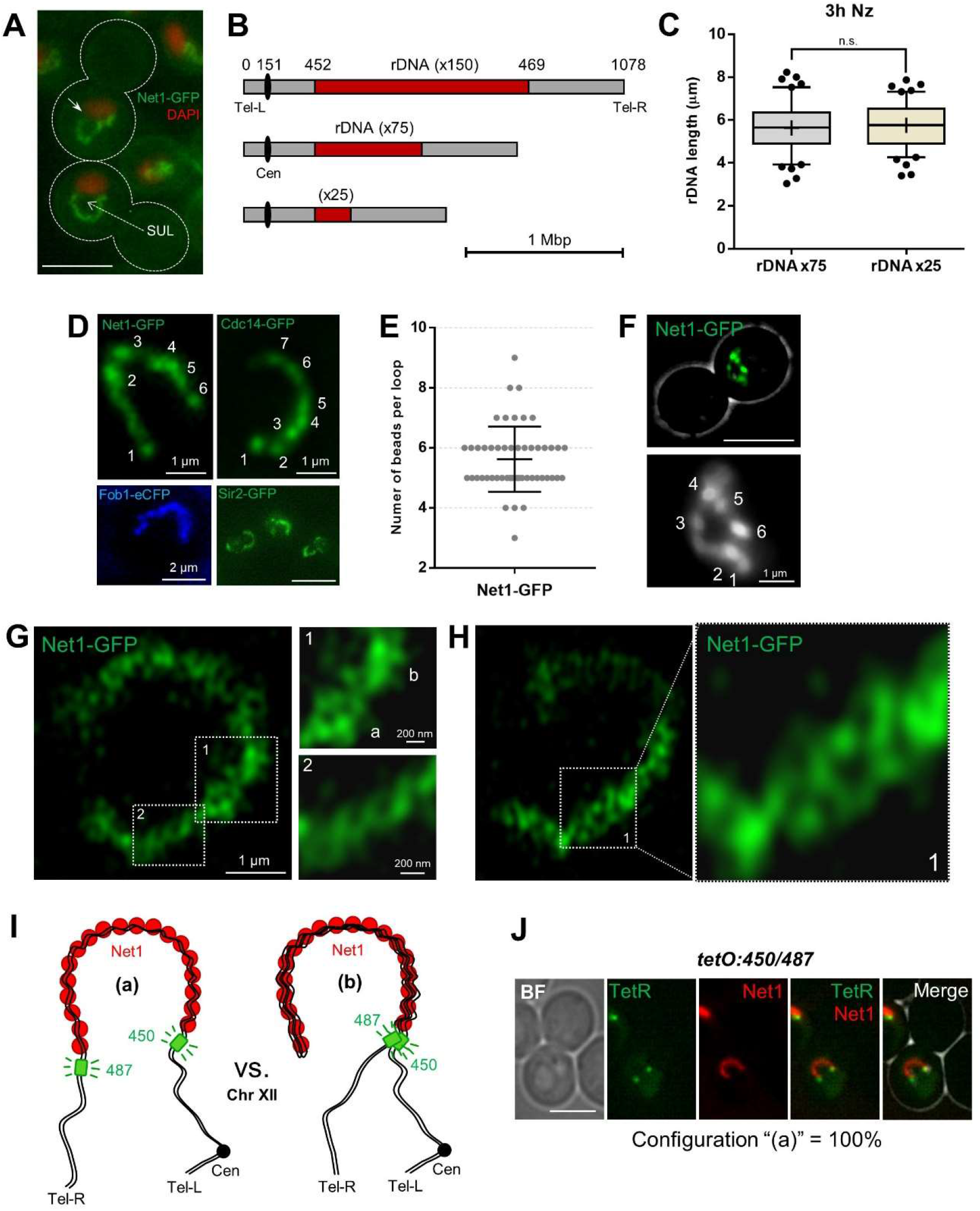
Fine structure of the mid-M rDNA loop. (**A**) Prototypical examples of the rDNA loop in the mid-M (nocodazole) arrest. The rDNA is labelled by Net1-GFP and the main nuclear mass by DAPI. The dotted arrow points to the space under the loop (SUL). The white arrow on the upper cell points to a small gap observed sometimes on one flank of the loop. (**B**) Scale drawing of chromosome XII with rDNA arrays bearing different numbers of its basic repetitive unit. The numbers above indicate Saccharomyces Genome Database (SGD) coordinates. (**C**) Length of rDNA loops from arrays with 75 (n=100 cells) and 25 (n=100 cells) copies. Boxplot: Whiskers represent 5-95 percentile. Mean shown as ‘+’. Dots represent outliers. (**D**) The rDNA loop comprises domains with distinctly enriched markers. Examples of rDNA loops with four specific markers: Net1 (deconvolved z-stack), Cdc14 (deconvolved z-stack), Fob1 (single z plane) and Sir2 (single z plane); the latter also labels telomere clusters and heterochromatinized loci. Numbers indicate counted “beads”. (**E**) Counted beads for the Net1-GFP rDNA loop (n=50 cells). Central horizontal solid line represents the mean and outer horizontal solid lines represent one standard deviation (S.D.). (**F**) The loop and its beads as seen by TIRF. (**G**and **H**) 120 nm superresolution of the rDNA loop. Inlets “1” features a section with two tight (G) (“a” and “b” separated with a constriction) or loose (H) braided threads. Inlet “2” features a section with knotty and twisted threads. (**I**) Schematics of two possible and mutually exclusive spatial configuration of the rDNA loop: (a), the loop has two bases, each comprising one of the rDNA flanks; and (b) the loop folds back, having both flanks residing on a single base. The green boxes indicate the *tetO* arrays inserted at 450 (rDNA proximal flank) and 487 (rDNA distal flank). (**J**) The rDNA loop in a strain with both 450 and 487 *tetOs*, the TetR-YFP and the Net1-eCFP (pseudo-coloured in red). All loops presented the configuration (a). In microscopic images, the scale bar represents 5 μm unless stated otherwise.

### The size of the rDNA array does not determine the length of the mid-M rDNA loop

Whereas it is undisputed that the rDNA is highly-organized in mid-M, previous studies raised concerns about whether the loop is a condensed state of the locus, at least from a longitudinal point of view ^10,36^. We previously measured loop lengths in a Net1-GFP strain with ~150 copies of the 9.137 kbp rDNA unit and obtained mean values of ~5.5 μm ^12^. This translates into a relative length of ~250 kbp/μm [150·9.137/5.5]. Taking into account that the length of one bp of naked B-form DNA is 0.34 nm, the estimated axial compaction ratio of the rDNA loop results in ~85 [250·0.34]. In some instances, the loop reached ~9 μm [~152 kbp/μm; compaction ratio ~52]. Previously, the axial compaction of chromosome arms in mid-M was estimated to be ~140 ^9^. Hence, the rDNA loop actually appears less compacted than a normal chromosome arm.

We started this work by questioning whether the loop length could be shortened by reducing the number of units of the rDNA array. We analysed the rDNA length in two strains with the rDNA size fixed at around 75 and 25 copies (Figures 1B; S1A and S1B). This was achieved through the deletion of the *FOB1* gene, responsible for the change in size of the array ^37^. Both strains were blocked in mid-M and the length of their rDNA loops were measured. We found no differences in their lengths (Figures 1C and S1C). The length was also equivalent to what we stated in our previous work with a ~150 copies rDNA ^12^. This implies that the compaction ratio of the rDNA can be lowered to, at least, ~10 (a loop of 25 copies reached 7.9 μm [0.34·25·9.137/7.9]). Theoretically, this packing ratio is close to the 10 nm chromatin fiber, assuming that all the array was fully and periodically coated by nucleosomes ^38^.

We conclude that (i) the size of the rDNA array does not determine the length of the rDNA loop, at least in a window of 25-150 copies; and (ii), conversely, the loop length is more or less fixed, with the compaction of the rDNA adapting to it, stretching the array if needed.

### Superresolution of the rDNA loop shows that it is organized as twisting threads of unequal density

We noticed that the rDNA loop does not appear homogeneously stained with different specific rDNA binding proteins (Net1, Fob1, Cdc14 and Sir2) (Figure 1D). Z-stacking and deconvolution analysis showed segmented beads lengthwise the rDNA loop. We could discern up to nine of these beads or domains, with a median of five per loop (Figure 1E), which we could further confirmed by Total Internal Reflection Fluorescence Microscopy (TIRFM) (Figure 1F). These beads likely represent the previously reported hierarchical organization of functional domains within the rDNA array ^39,40^.

We next approached the fine characterization of the mid-M rDNA loop by adding visualization through confocal superresolution microscopy (CSM). CSM showed that the rDNA loop was ~200 nm thick and comprised twisting of up to two threads, with sections resembling a spring (Figures 1G and 1H). We could also distinguish regions of extensive knotting separated by constrictions (“a” and “b” in Figure 1G), which probably account for the beads observed under epifluorescence microscopy.

The temporal behavior of threads and knots was rather dynamic, although the overall length and shape of the rDNA loop turned out to be quite stable over the course of short time-lapse movies (Movie 1).

Altogether, we pinpoint that the loop is not homogeneously compacted and resembles a spring. Both features may explain why the loop retains a constant length upon reducing the locus size.

### The bases of the rDNA loop are the flanking sequences of the rDNA array

We reasoned that the two twisting threads observed in the loop by CSM may comprise either the cohesed sister chromatids, resolved at 120 nm laterally, or the two halves of a coiled coil rDNA array (Figure 1I; “a” and “b”, respectively). In support of the latter (“b”), we often found small gaps between one edge of the Net1 signal and the DAPI (Figure 1A, arrow). In this scenario, the low compaction ratios calculated above ought to be further divided by two. In order to differentiate between these two models, we combined into a single Net1-eCFP strain two bacterial *tetOs* arrays that we have previously designed to flank the rDNA array; *tetO:450* (rDNA proximal flank) and *tetO:487* (rDNA distal flank). This strain also carries the *tetO*-specific binding protein TetR-YFP. If the rDNA loop comprised an extended array that coils back (“b”), we should see both *tetOs* localizing at the same loop edge. On the contrary, if the rDNA loop only comprised the two sister chromatids travelling all along its length (“a”), we should see each *tetO* at each edge of the loop. We observed the “a” configuration in all cases (Figure 1J). This flanking positioning is in agreement with previous FISH data for the rDNA borders at the chromosome XII right arm (cXIIr) ^41^. This configuration also points out that the two threads observed by CSM must correspond to the sister rDNA chromatids.

Similar configuration results were observed when we employed a partner comprising *tetO:450* and a *tetO:1061*, which localizes near the cXIIr telomere. Even though the *tetO:1061* showed a looser localization, it was often close to the opposite *tetO:450* base (Figure S2A). The distance between the *tetO:1061* and the distal rDNA flank ranged from 0.2 to 4 μm (Figure S2B). In short time-lapse movies, the *tetO:450* remained immobile and attached to the loop base, while the *tetO:1061* moved rapidly and extensively (Movie 2). Shorter distances to the nearest rDNA flank were measured for the *tetO:194*, which settles next to the chromosome XII centromere (Figure S2B). However, relative distances (μm/kbp) were the opposite, taking into account that the *tetO:194* is ~200 Kbs from the proximal rDNA flank, while the *tetO:1061* is ~600 Kbs away from the distal rDNA flank (mean relative distances of 0.0025 and 0.0017 μm/kbp, respectively). If we assume an unlikely straight line between the telomere/centromere and the corresponding rDNA flank, the resulting mean apparent compaction ratio would be 205 for the centromere-proximal flank tract, and 280 for the distal rDNA flank-telomere tract (Figure S2C). In order to fit with the compaction ratio of 140, both tracts are probably curly or eventually fold back, especially the distal rDNA flank-telomere tract (Figure S2D).

We conclude that the bases of the rDNA loop are the flanking rDNA regions and that the rest of the chromosome XII right arm usually locates out of the loop.

### The space under the rDNA loop is not nuclear and is occupied by the vacuole

The second remarkable feature of the mid-M arrest is the SUL, the large space between the loop and the main DAPI-stained nuclear mass. To our benefit, in the previous strains, not all TetR-YFP molecules bind to the *tetOs*. A pool of TetR-YFP freely circulates within the nucleus, labelling the nucleoplasm and, hence, marking the shape of the nucleus. When observing the Net1-eCFP loops, the free TetR-YFP surrounded the rDNA in all cases. Strikingly, the SUL was void of TetR-YFP, especially in those loops which were large and highly bended (horseshoe-like) (Figure 2A). We hypothesized that this space may comprise other parts of the nucleolus, which could be somehow free of the TetR, perhaps through liquid-liquid phase separation ^24,42^. However, we found neither rRNA-processing proteins (Figure 2B) nor RNA (Figure 2C). In both cases, the mid-M arrest made these markers adopt a horseshoe-like loop as well. Alternatively, the SUL could be filled up by part of the nucleus devoid of freely circulating proteins; for instance, heterochromatinized DNA. However, we did not find histone-coated (Hta2-mCherry) DNA, DAPI (after overexposing) or ultrasensitive YOYO-1 DNA stain within that space (Figures 2D and 2E).

**Figure 2.**
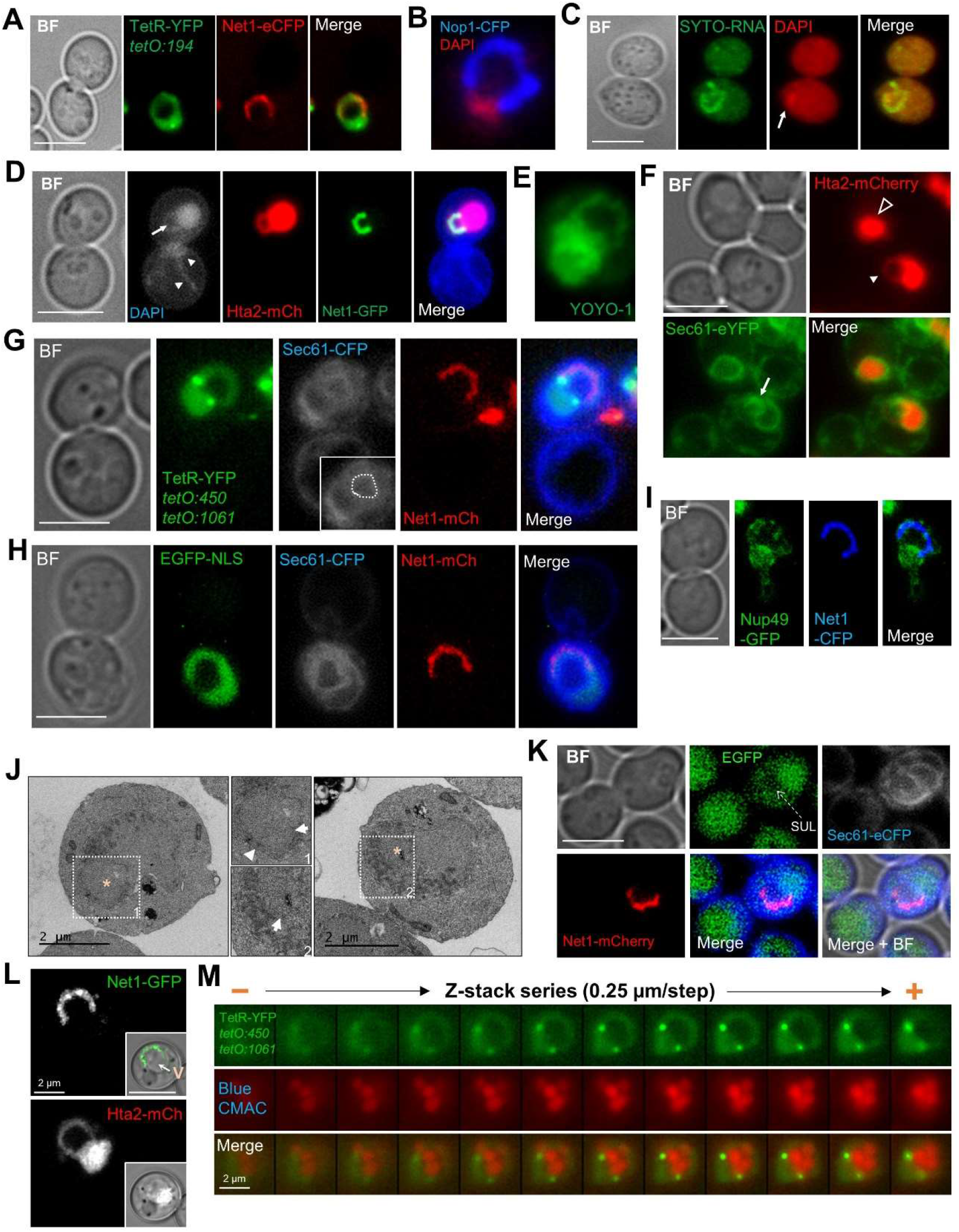
The space between the rDNA loop and the rest of the nuclear mass is not nuclear and is occupied by the vacuole. (**A**) The heterologous TetR that freely circulates in the nucleoplasm does not label the SUL. (**B**) The nucleolar SSU rRNA processome protein Nop1 also forms a horseshoe-like loop. (**C**) The RNA marker SYTO-RNA, which strongly labels rRNAs, also forms a horseshoe loop. (**D**) Overexposed DAPI (shown in grey) and fluorescently-tagged histone Hta2 (H2A) co-label the rDNA loop (Net1-GFP), which is seen as a handle in a handbag-like nucleus. The arrow points to the handle, whereas arrowheads point to non-nuclear (mitochondrial) DNA. (**E**) The ultrasensitive dsDNA stain YOYO-1 also stains a handbag-like nucleus in mid-M. (**F-J**) The SUL is surrounded by nuclear membranes. (**F**) The space within the Hta2 handle is surrounded by the nER membrane marker Sec61. Two cells are represented, one with (filled arrowhead) and one without (hollow arrowhead) the histone handle. (**G**) Sec61 delimits both the internal and external boundaries of the nuclear space (the TetR handle) that contains the rDNA loop (Net1-mCherry). The Sec61-eCFP channel is shown in grey. The internal (handle-SUL) Sec61 signal is highlighted with a dotted drawing in the inlet image. (**H**) Like in G but with the nucleoplasm labelled with an EGFP-NLS construct instead of the TetR. (**I**) The rDNA loop (Net1-CFP) is also surrounded by the nuclear pore complex component Nup49. (**J**) TEM images of mid-M arrested cells with a nuclear morphology compatible with the ones shown in F-I. The dotted squares mark areas of interest, shown in more detail in the middle. Electrodense material represents the nucleolus. SULs are indicated by asterisks. The arrow and the arrowhead point to the internal (handle-SUL) and the external (handle-cytosol) double membrane, respectively. (**K**) Cytosolic EGFP weakly stains the SUL. The strain also bears Sec61-eCFP and Net1-mCherry to delimit the SUL (pointed with the dotted arrow). (**L**and **M**) The vacuole resides in the SUL. (**L**) The rDNA loop (Net1-GFP and Hta2-mCherry handle) partly surrounds the vacuole (V), pointed with the arrow in the inlet DIC image. Photos obtained through CSM. (**M**) Montage of z stack images of a TetR handbag-like nucleus where the vacuole lumen has been stained with Blue CMAC. Scale bars represent 5 μm unless stated otherwise.

In light of these negative labelling, we next questioned whether the SUL was nuclear. For addressing this, we added a third fluorescent marker to the previous set of strains, the endoplasmic reticulum membrane translocator Sec61, which labels the perinuclear endoplasmic reticulum (nER), this being a continuum with the outer nuclear membrane. We found that the Net1 loop was surrounded both externally and internally by Sec61 (Figures 2F and 2G). A similar pattern was obtained when an EGFP-NLS replaced TetR-YFP as the nucleoplasm reporter (Figure 2H). This was further confirmed by a nuclear envelope (NE) marker, the nuclear pore complex protein Nup49. Nup49 is more specific for the NE than Sec61 but its labelling is punctated ^43^. Nonetheless, we could confirm that Nup49 labels the bases, as well as the internal face, of the loop (Figures 2I and S3A). In order to get further support for the claim that the SUL is not nuclear, we performed transmission electron microscopy (TEM). We found examples where the nucleus appears as a handbag, with a thin handle surrounded by a double membrane, and with the electrodense nucleolus close to or within this handle (Figure 2J).

After realizing that the SUL was not nuclear, we added a cytosolic EGFP to a Sec61-eCFP Net1-mCherry strain to confirm the SUL was cytosolic. To our surprise, the EGFP signal was weaker in the SUL (Figure 2K). A key hint to understand the nature of the SUL came from the visual comparison of the Net1 loop and the Hta2 handle with the entire mid-M cell as seen through the transmission light (bright field). We noticed that most large horseshoe-like loops appear to surround the vacuole (Figure 2L). In addition, other loops also appear to transverse vacuoles (Figures S3B and S3C). Thus, we checked whether the vacuole was occupying the SUL. We first employed the vacuolar membrane stain FM4-64. For reasons we do not know yet (also confirmed by M. Moriel-Carretero, personal communication), the FM4-64 staining pattern was weak, diffuse and dotted on those horseshoe-like loop coincident with the vacuole (Figure S3C). We thus employed instead the vacuolar lumen stain Blue CMAC, finding that it strongly stained the SUL. Z-stack imaging, deconvolution, fluorescence intensity profiles and orthogonal projections confirm that the SUL is occupied by the vacuole (Figures 2M and S3D-G).

### The rDNA horseshoe loop coexists with protruding rDNA bars that bend around vacuoles

In the following chapters, we focused on the origin of the horseshoe rDNA loop with the vacuole in the SUL. An important hint came from observations that slightly differ from the morphological patterns described above.

Despite the horseshoe rDNA loop is the most remarkable morphology of the rDNA array in mid-M, other bar-like morphologies are observed as well. A close look at the Net1-GFP Hta2-mCherry strain showed that Net1-GFP horseshoe loops represented ~20% of the Nz-blocked cells, ~35% presented a protruding bar-like Net1-GFP signal, and another 20% showed a Net1 bar that was either attached to or crossed through the main nuclear mass, with no SUL (Figures 3A-D). The remaining non-loop morphologies, oval and resolved (two separated Net1 cluster in a single nucleus), were a minority and seemingly correspond to cells that have not reached or have passed the mid-M arrest, respectively.

**Figure 3.**
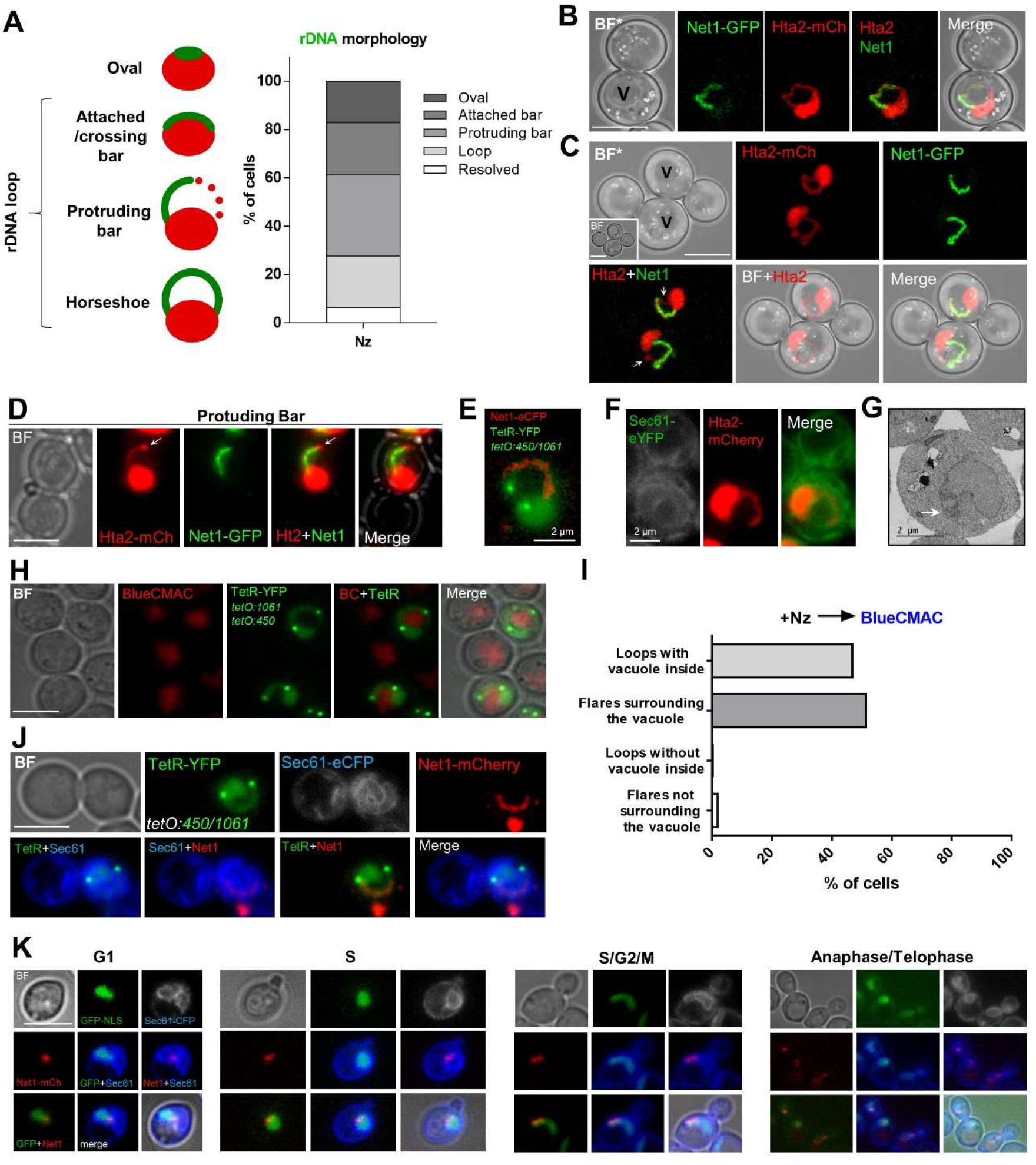
The rDNA loop arises when bars of rDNA protruding out of the main nuclear mass wrap around the vacuole. (**A**) Quantification of rDNA morphologies (Net1-GFP) relative to the main nuclear mass (Hta2-mCherry). (n=217 cells). (**B**) Example of a full loop (Net1 loop and Hta2 handle overlaps almost completely) as seen by CSM. BF*, extended depth of focus of the bright field channel. (**C**) Examples of a protruding rDNA bar in the context of an Hta2 handle (Net1 bars and Hta2 handles overlap partly only), as seen by CSM. The arrows point to the Net1/Hta2 gaps. The inlet BF image shows a single z plane where vacuoles can be easily identified. (**D**) Example of a protruding rDNA bar in the context of a protruding Hta2 bar. The arrows point to the denser Hta2 signal ahead of the Net1 tip. (**E**) The telomeric flank of the rDNA localizes ahead of the protruding rDNA bar. (**F**) The protruding rDNA bar is surrounded by the nER, which forms a flare. (**G**) TEM of a NE flare (pointed by an arrow) with the electrodense nucleolus inside. (**H**) Examples of nuclear flares surrounding the vacuole, whose lumen is stained with Blue CMAC (pseudo-coloured in red). (**I**) Quantification of loops and flares surrounding the vacuole (n=109 cells). (**J**) Example of the closest probable precursor of the rDNA loop originating from a protruding rDNA bar that wraps around a vacuole. Note the snail shell morphology of the NE. (**K**) The reported mid-M arrest morphologies are not present during a normal cell cycle.

The protruding bars appear to be anchored to the bulk of the nuclear mass through one base only (Figure 3C and 3D). We observed two classes of these protruding bars. In the first class (about two third of the cases), the rDNA overlaps partly with an Hta2 handle, suggesting that flanking regions of the rDNA can belong to a wider vision of horseshoe loops (Figure 3C). Single-cell time lapse of these two presentations of the Hta2 handle show they are interchangeable (Movie 3; e.g., nuclei 2, 4, 6 and 13). These, together with the fact that attached/crossed bars likely represent top views of handles (Movie 3, nuclei 3 and 15; Movie4, nuclei 3-5), imply that horseshoe loops are more frequent than the observed 20% at first glance. In the second class of protruding rDNA bars (10% of all morphologies), the Hta2 also forms a protruding bar, almost fully coincident with the Net1-GFP signal (Figure 3D). In this class, the tip of the rDNA protruding bar often contained a brighter Hta2 spot (arrow). This spot must correspond to a hypercondensed state of the 600 Kbs of the distal cXIIr. In agreement with this, data from the *tetO:450 tetO:1061* TetR-YFP Net1-eCFP strain showed that one *tetO* was found at the tip of nuclear/Net1 morphologies compatible with this class of rDNA bars (Figure 3E). Moreover, these rDNA bars protrude out of the main nucleoplasmic region into a bulge of TetR-YFP signal (Figure 3E), and is surrounded by the NE (Figure 3F). This configuration of having the nucleolus into nuclear bulges was also confirmed by TEM (Figure 3G). These results are in full agreement with the nucleolus-containing NE “flare” previously described by the Cohen-Fix’s lab ^32,33^.

Remarkably, rDNA bars, and their surrounding constituents (nucleoplasm and NE flares), showed extensive bending, which were clearly reminiscent of incomplete states of the horseshoe loop (Figures 3E and S4A). Hence, we tested whether the vacuole was also associated to rDNA bars/flares. We found it was (Figures 3H and 3I). Not only that, it also appears that the vacuole drives the bending of the protruding bars until the bar becomes a horseshoe loop (Figure 3H and 3J). In addition, several of these examples suggest that the horseshoe loop may comprise a rather extended, and sometimes bilobed, nucleus that bends onto itself until acquiring the frontal appearance of a unilobed normal nucleus (Figures 3J; S4B and S4C; Movie 3, nucleus 15). None of these remarkable morphologies were seen in asynchronous cultures (Figure 3K), although the nucleus may appear squeezed between the vacuole and the plasma membrane in some cells, as if the NE is a malleable body which must seek allocation between two other stiff bodies, the vacuole and the cell wall (Figure 3K, S/G2/M example; S4D and S4E).

### The horseshoe rDNA loop that wraps around the vacuole requires the absence of microtubules

Previous works demonstrated that the flare morphology is characteristic of cells blocked in mid-M ^33^. Nz is the most common experimental tool to achieve the mid-M arrest. Nz depolymerizes microtubules, which dismantles the spindle apparatus and activates the spindle assembly checkpoint (SAC) ^44,45^. With an active SAC, the anaphase-promoting complex (APC) activator Cdc20 is tightly bound and inhibited by the SAC components Mad2 and Mad3 ^46,47^. Considering the effect of Nz on microtubules and the SAC, we next chose to study the effect of arresting cells in mid-M by other means. We planned two different strategies that preserved the spindle, yet they differ in the activation mode of the SAC (Figure 4A). On the one hand, we depleted Cdc16, an essential component of the APC, by creating a *cdc16:aid* conditional allele. Degradation of Cdc16-aid can be conditionally triggered by adding the auxin indole-3 acetic acid (IAA) to the medium (Figure S5A, B) ^48^. Under this condition, the APC can be inactivated without interfering with the spindle and/or the SAC, although there are reports about a mimicking or possible activation of the SAC upon APC inactivation ^49^. On the other hand, we made use of another strain carrying a *P_GAL1_*-*MAD2-MAD3* construction, which overexpresses a fusion protein of these two key SAC players, yielding an active SAC in galactose. Under this condition, the active SAC maintains the mid-M arrest by keeping the APC inactive^50,51^.

**Figure 4.**
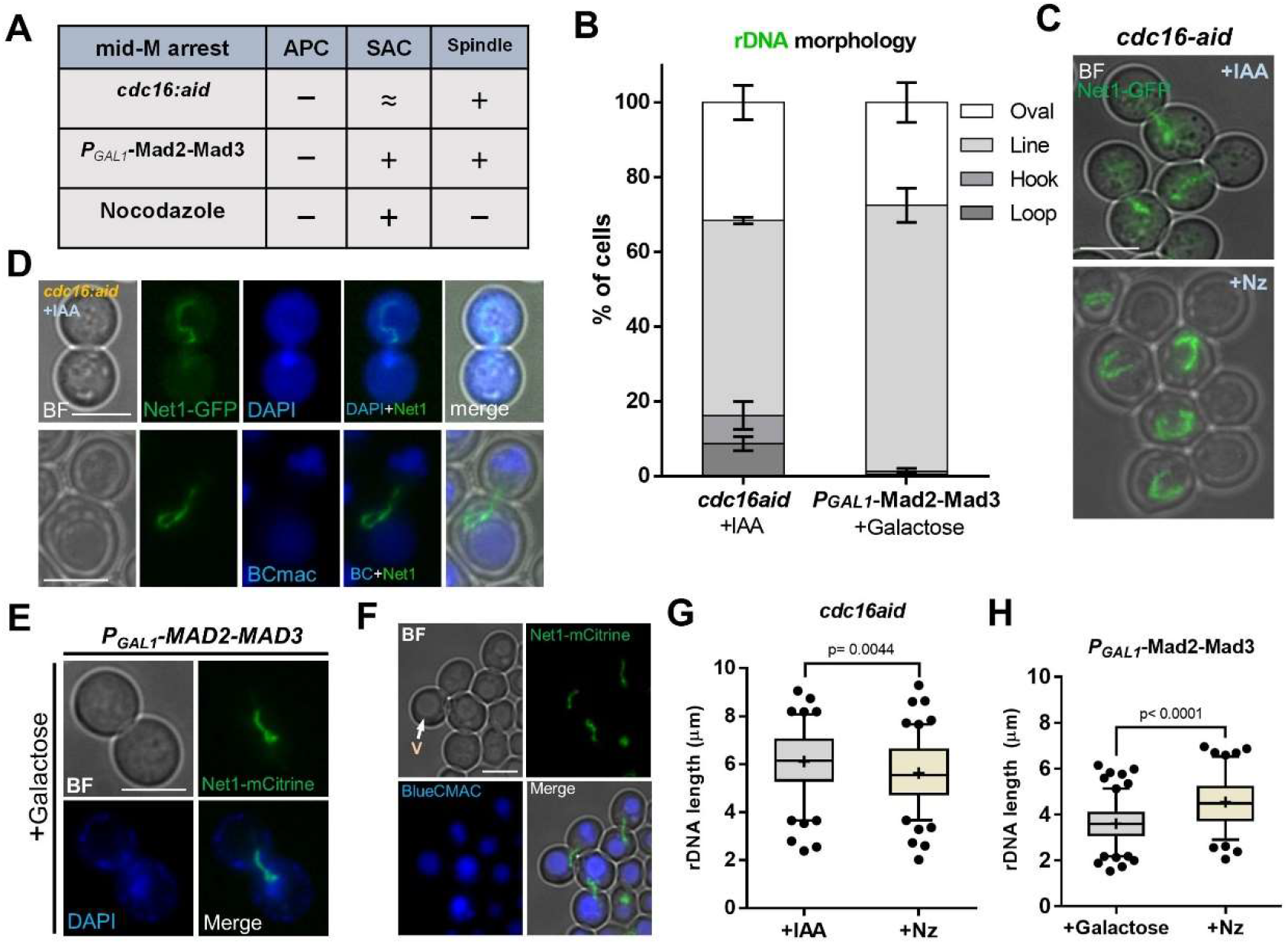
The rDNA loop in mid-M arrests that preserve the spindle. (**A**) Summary of the differences between three mid-M arrests (depletion of Cdc16, overexpression of the Mad2-Mad3 chimera, and Nz). APC, anaphase-promoting complex; SAC, spindle assembly checkpoint. The plus sign indicates either active (APC, SAC) or present (spindle); the minus sign indicates the opposite; “≈” indicates mimicking or possible activation. (**B**) Morphology of the rDNA after Cdc16-aid depletion and Mad2-Mad3 overexpression (mean ± S.E.M; n=3 independent experiments (>100 cells each)). (**C**) Representative micrograph of the rDNA lines observed after depleting Cdc16-aid (upper picture). The same strain under Nz alone is shown underneath for comparison. (**D**) Representative micrographs of the rDNA hooks observed after depleting Cdc16-aid. The upper picture shows that the rDNA and the gross DNA mass (DAPI) reside in different cell bodies. The lower picture shows a hook relative to its nearest vacuole (BC staining). (**E-F**) Representative micrographs of the rDNA lines observed after overexpressing Mad2-Mad3. (**E**) Net1 together with DAPI staining. (**F**) Net1 together with BC staining. (**G** and **H**) Length of the rDNA (Net1) in previous mid-M arrests. (**G**) Cdc16-aid plus IAA (>100 cells) vs. Cdc16-aid plus Nz (>100 cells). (**H**) *PGAL1-MAD2-MAD3* in YPgalactose (>100 cells) vs. YPD plus Nz (>100 cells). Boxplot: Whiskers represent 5-95 percentile. Mean shown as ‘+’. Dots represent outliers. In micrographs, the scale bar represents 5 μm; BF, bright field; V, vacuole; BC, Blue CMAC

We observed a general pattern that was shared by both non-Nz mid-M strategies, and that greatly differed from Nz-arrested cells. Although an organized bar-like rDNA (Net1-GFP) was seen in all conditions, the horseshoe loop was largely absent and, instead, a straight bar (line) often orientated in the chromosome division axis (polar axis) was the major outcome (Figure 4B-F). This shift in the rDNA loop morphology was not a consequence of the newly introduced alleles, as Nz still leads to horseshoe loops in these strains (Figure 4C and S5C). The bending of the bar was less prominent than in Nz, yet it sometimes acquired a hook shaped morphology (Figure 4D). The bar bending and the hook still follow the vacuole in many instances (Figure 4D, F). Interestingly, the rDNA length was slightly larger in the straight line than in the horseshoe loop, pointing out that the rDNA is even more stretched in such configuration (Figure 4G). However, in galactose, a condition needed to active the SAC in the *P_GAL1_*-*MAD2-MAD3* strain, the line was shorter (Figure 4H).

We conclude that the absence of microtubules is a prerequisite to acquire the horseshoe rDNA loop that wraps around the vacuole. However, the mid-M arrest is sufficient to change the rDNA morphology into the highly organized bar.

### The rDNA bar and the nuclear flare depend on active TORC1 and membrane phospholipid synthesis

We have previously shown that conditions that inactivate the TORC1 complex dismantle the rDNA loop, including mild acute HS ^12^. Intriguingly, many reports have demonstrated that TORC1 inactivation activates autophagy, including nucleophagy, thereby promoting the nuclear-vacuolar interaction ^52,53^. According to the results shown above, it appears counterintuitive that the horseshoe rDNA loop is absent when the influence of the vacuole on the nucleus should be maximum. To get further insights, we studied the rDNA and nuclear mass morphologies in cells transiting through stationary phase, when TORC1 activity is expected to be low, autophagy high, and most cells appear swollen and with a large vacuole compressing the rest of the cell organelles. In this condition, the rDNA (Net1-GFP) was hyper-compacted, and its structure was barely modified by the vacuole (Figure 5A). Moreover, we could not observe any histone (Hta2-mCherry) handles. The addition of Nz did not change this pattern, demonstrating that Nz only elicits its effects on the rDNA in growing cells, when Nz lead to the mid-M arrest.

**Figure 5.**
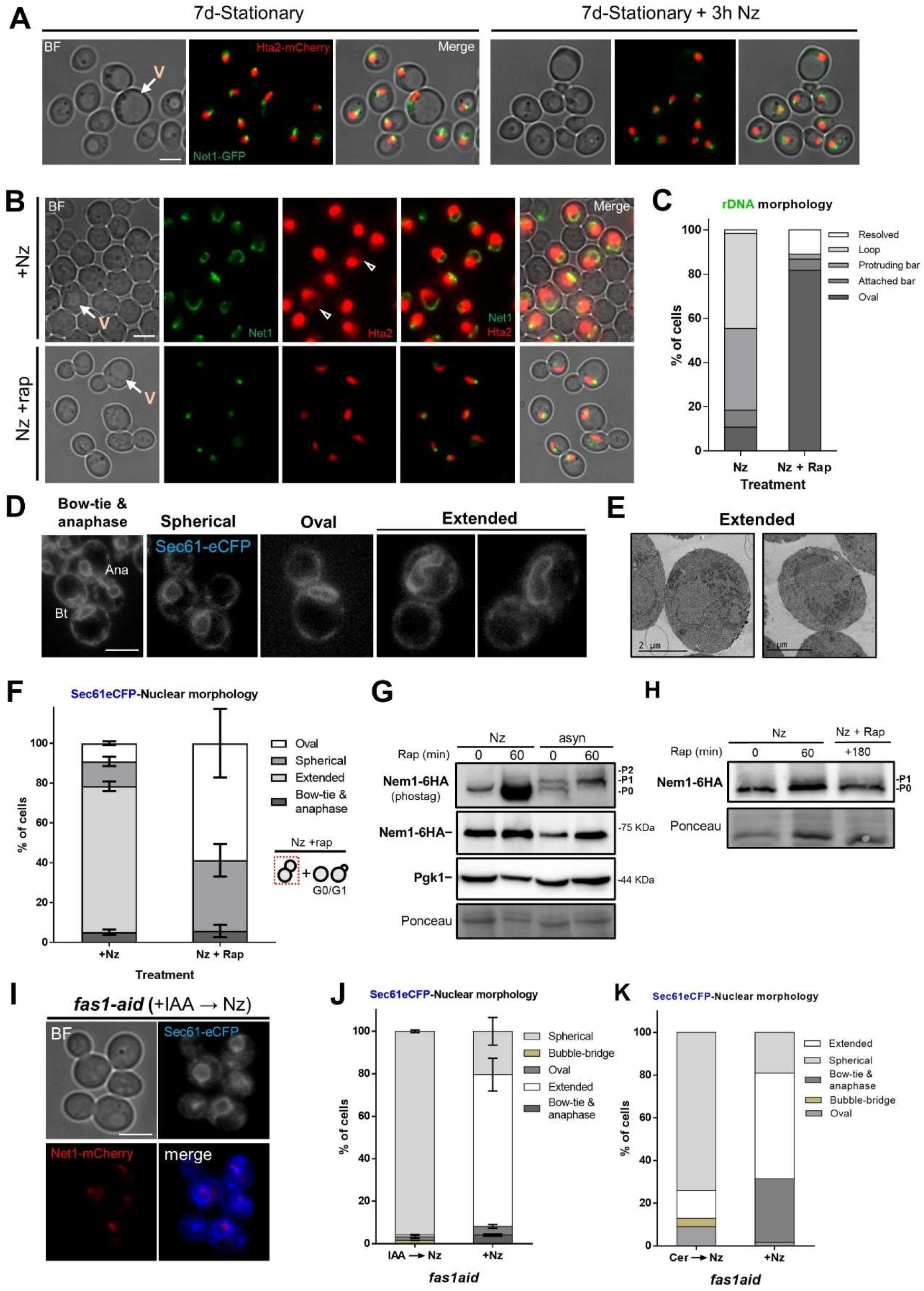
The rDNA loop requires TORC1 and membrane synthesis. (**A**) The rDNA loop is not present in stationary phase (7 days of continuous growth in a flask) upon Nz. (**B-C**) Rapamycin prevents rDNA loop formation in Nz-treated cells. (**B**) Representative cells upon Nz alone or Nz plus rapamycin; arrowheads point to Hta2 handles. (**C**) Quantification of rDNA morphologies. Categories based on Net1 and Hta2 comparisons as in Figure 3A. (**D**) Representative micrographs of nuclear morphologies seen by Sec61 in mid-M arrests. Bt, bow-tie nucleus; Ana, dumbbell shaped anaphase nucleus (the cell has escaped from the arrest). Extended is a mid-M specific category that acquire finger-like projections and can take several forms (two examples shown). (**E**) TEM micrographs of mid-M arrested cells with a nuclear morphology compatible with the extended category. (**F**) Morphology of the NE (Sec61) in Nz and Nz plus rapamycin (mean ± S.E.M; n=3 independent experiments (only mid-M cells counted; >100 cells in Nz, >60 cells in Nz + rap). (**G**) Western blot of Nem1-6HA under Nz alone and Nz followed by rapamycin. An equivalent rapamycin regime over an asynchronous culture was performed as well for reference. The position of the three expected bands is indicated in the electrophoresis run with Phos-tag (top image). The others were standard SDS-PAGEs controls. (**H**) An independent repetition of part of G, which also includes a concurrent Nz rapamycin treatment. The protein electrophoresis was with Phos-tag. (**I-J**) Depletion of the fatty acid synthetase beta subunit Fas1 prevents nuclear extension upon Nz treatment. (**I**) Representative micrographs. (**J**) Morphology of the nER (Sec61) in Nz and IAA→Nz (mean ± S.E.M; n=2 independent experiments (>190 cells each; only mid-M cells counted)). (**K**) Inhibition of fatty acid synthetase with cerulenin also prevents nuclear extension upon Nz treatment. The NE morphology was quantified as in J (>100 cells). In micrographs, the scale bar represents 5 μm unless stated otherwise; BF, bright field; V, vacuole.

Next, we studied the effect of rapamycin addition, a well-known inhibitor of the TORC1 complex ^54–56^. For this, we first compared a Nz arrest to a concomitant Nz plus rapamycin treatment. On the latter, we found that the rDNA morphology was mainly oval without histone handles, whereas the main morphologies in Nz alone were the expected rDNA loops with either histone handles or protruding bars (Figures 5B and 5C). Next, we added rapamycin to the cells previously arrested in Nz; after one hour of rapamycin treatment, these structures were dismantled into compact ovals, short lines or hyper mini-loops/handles of Net1/Hta2 (Figure S5D), as we have shown before ^12^.

On previous sections, we show that horseshoe loops and bars are associated with extended nuclei, as seen by rDNA, histone and NE markers. Thus, we studied nuclear morphology under Nz treatment and TORC1 inhibition by following the NE shape with Sec61-eCFP. In growing cells, most nuclei appear either spherical or oval shaped (Figures 5D and S5E), except in those cells transiting M phase where the nucleus is stretched along the mother-daughter axis. In such cases, two morphologies are distinguished. The first one is bow-tie shaped, which is shared by cells that are in late metaphase and early anaphase ^57^. The second one is dumbbell shaped and corresponds to cells in late anaphase. Upon Nz treatment, the addition of the NE flare made nuclei also appear extended and/or stretched but not in the mother-daughter axis (Figures 5D and 5E; S5E). However, under Nz plus rapamycin the nuclear morphology was mainly spherical/oval again (Figures 5D and F). Similarly, cells in stationary phase presented a spherical/oval morphology and, once again, the addition of Nz did not change this pattern (Figure S5F).

In a mid-M block phospholipid synthesis is unabated, and the nuclear membrane expands around the region that contains the nucleolus ^32,33^. Several lines of evidence pinpoint the Nem1-Spo7/Pah1 complex as a central player for the control of nuclear membrane expansion. This complex is involved in the balance between membrane phospholipids during growth conditions and lipid droplets during starving conditions; an active Nem1-Spo7/Pah1 complex shifts the balance towards the latter ^30,58^. The TORC1 complex regulates the activity of Nem1-Spo7/Pah1 by keeping Nem1 unphosphorylated and inactive, so that the phospholipid synthesis is favored ^59^. Accordingly, Nem1-Spo7/Pah1 mutants display flares and extensions in growing cells, similar to those presented herein under Nz ^31,32^. For this reason, we studied the phosphorylation status of Nem1 under Nz treatment and TORC1 inhibition. We arrested cells in either Nz alone or Nz followed by rapamycin addition for 1h, and further compared these conditions to rapamycin treatment in asynchronous cultures (Figures 5G and S5G). We observed the same pattern of major phosphorylation shifts that have been described before in asynchronous exponentially growing cells: two bands, P0 and P1, and a third band, P2, after rapamycin addition ^59^. Strikingly, only the P0 band was seen in Nz. This un(hypo)phosphorylated state suggests a strong Nem1 inhibition in Nz, which was modified only modestly by rapamycin, either after or concomitant to Nz addition (Figure 5H).

We further decided to investigate the role of lipid metabolism on the phenotypes described here. It has been shown that activation of mRNAs encoding lipogenic enzymes (Acc1, Fas1 and Fas2), all involved in fatty acid synthesis, increased in G_2_/M ^60^. Previous studies have also shown that biosynthesis of fatty acids is necessary for the extension of the nuclear membrane ^33,61–63^. Thus, we drew our attention to fatty acid synthesis and its relation to nuclear membrane growth in Nz. We made a Fas1-aid chimera and tested both NE (Sec61-eCFP) and rDNA (Net1-mCherry) morphologies when Fas1 was degraded before Nz addition (Figure 5I and 5J; S5H-J). Degradation of Fas1-aid in IAA was partial (~66% drop in protein levels; Figure S5H); however, this was sufficient to prevent NE extensions in Nz (Figure 5I and 5J; S5I). In addition, we corroborated these findings by employing cerulenin, a specific inhibitor of fatty acid biosynthesis ^64^. We treated cells with cerulenin, one hour before the addition of Nz, which also resulted in spherical nuclei (Sec61-eCFP) and compacted Net1-mCherry signals (Figure 5K). Interestingly, we also observed more spherical NE morphologies within the bow-tie subgroup (Figure 5J and 5K; S5J; “bubble bridge”), suggesting a stiffer NE when fatty acid biosynthesis is inhibited.

### The rDNA loop does not depend on known nuclear-vacuolar interactions

From our previous results, the shape of the malleable NE appears to be highly influenced by the stiffer vacuole. When the NE becomes enlarged in mid-M blocks, the vacuole serves as a template on which the extended NE bends around. In this context, the length of the rDNA in protruding bars and horseshoe loops may depend on how intimate the nuclear-vacuole relationship is. For this reason, we decided to study the role of the nucleus-vacuole junctions (NVJs) in the morphology of the rDNA. The NVJs are formed through the formation of Velcro-like interactions between the vacuolar protein Vac8 and the outer nuclear membrane (continuum with the nER) protein Nvj1, which mediate piecemeal microautophagy of the nucleus ^65,66^. Similarly, nER-vacuole contacts are established as sites for lipid droplet biogenesis, which include the proteins Nvj1, Mdm1, Nvj3, Nvj2 and Vac8 ^67,68^. We tagged the Net1-GFP in wild type, double (*mdm1Δ*/*nvj3Δ*) and quadruple (*mdm1Δ*, *nvj1Δ*, *nvj2Δ* and *nvj3Δ*) “ΔNVJ” mutants, of which, ΔNVJ is known to increase the nER–vacuole inter-organelle distance ^67^. Cells were arrested in Nz for 3h and the rDNA structure visualized as before. Surprisingly, we found that the majority of the cells presented an rDNA loop, even in the quadruple ΔNVJ mutant (Figure 6A and 6B). Moreover, when we stained the cells with BlueCMAC in the ΔNVJ mutant, the loops still wrap the vacuole (Figure 6C). We conclude that nuclear/(nER)-vacuole contacts, are not a prerequisite for the formation of the horseshoe rDNA loop.

**Figure 6.**
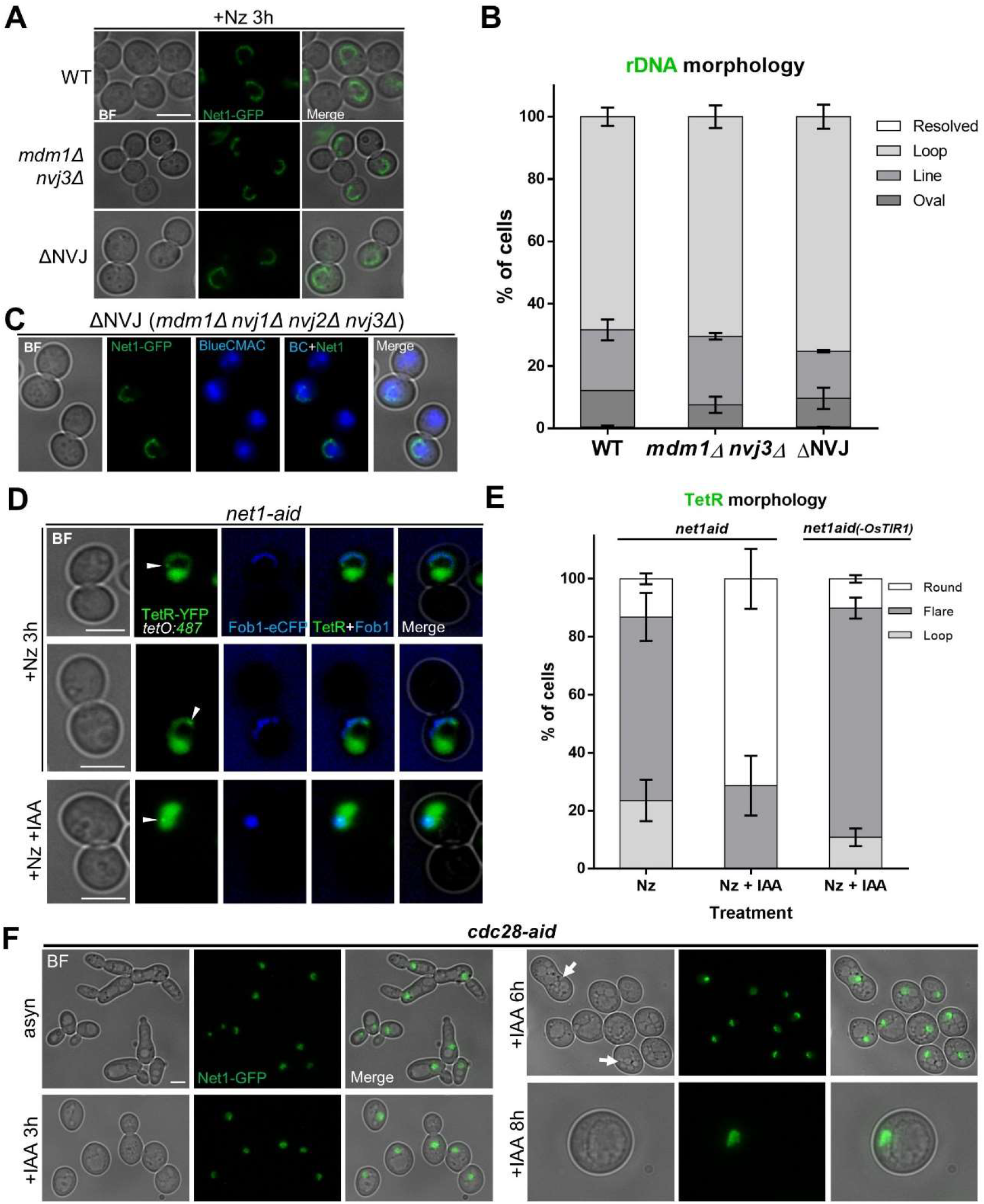
The rDNA loop is independent of nucleus-vacuole junctions and requires high CDK and an inactive Cdc14. (**A-C**) The mid-M arrest with Nz forms horseshoe rDNA loops in mutants for nucleus-vacuole junctions. (**A**) Representative mid-M cells in the WT and mutants that affect nucleus-vacuole interactions at different levels. (**B**) Quantification of rDNA morphologies upon Nz in the different mutants (mean ± S.E.M; n=2 independent experiments (>200 cells each)). (**C**) Representative mid-M cells in the ΔNVJ quadruple mutant. The rDNA (Net1) and vacuoles are shown. (**D-E**) The ectopic release of Cdc14 (by depletion of Net1-aid, the Cdc14 inhibitor) prevents the formation of the rDNA loop. (**D**) Representative mid-M cells in Nz alone and Nz plus IAA. The arrowhead points to the *tetO:487*. (**E**) Quantification of rDNA morphologies under Nz alone and Nz plus IAA (mean ± S.E.M; n=3 independent experiments (>100 cells each)). A Nz plus IAA in a *net1-aid* strain without the auxin-mediated degron system (OsTIR1) is included as well. (**F**) The cyclin dependent kinase Cdc28 is required for the rDNA loop. Cdc28 was depleted during several hours from an asynchronous culture. Cells became arrested in G1, enlarged and with fragmented vacuoles (arrows) but the rDNA did not shift into a loop. In micrographs, the scale bar represents 5 μm; BF, bright field; V, vacuole; BC, Blue CMAC

### The rDNA loop is not formed when cells are not biochemically in G2/M

Biochemically, the mid-M is characterized by high cyclin dependent kinase (CDK/Cdc28) activity, as well as an inactive Cdc14 ^69^. Cdc28 and Cdc14 are the master cell cycle kinase and phosphatase, respectively. These biochemical features are shared within a wider cell cycle window, which includes S and G2/M. Thus, we determined the rDNA morphology in conditions that differ from this S/G2/M biochemistry.

Firstly, we studied the unscheduled ectopic release of the Cdc14 phosphatase under Nz treatment. Cdc14 phosphatase is kept inactive and bound to its inhibitor Net1 in the nucleolus for most of the cell cycle, until its nucleolar release on early anaphase ^70,71^. Upon anaphase onset, Cdc14 inhibits transcription by RNA polymerase I on rDNA, which allows condensin to compact the rDNA ^72^. We constructed a strain carrying a *net1-aid* conditional allele, instead of using temperature sensitive alleles that interfere with the rDNA loop through the HS response ^11,12^. This strain also carried the TetR-YFP/*tetO:487* and Fob1-eCFP, which Net1 binds to, to visualize the rDNA and its distal flank. Degradation of Net1-aid in IAA (Figures S6A and S6B), ought to allow the ectopic release of Cdc14 outside anaphase, as reported for the *net1-1* allele ^71^. Cells were arrested in Nz or Nz plus IAA and the different rDNA morphologies scrutinized. We found bended bars and horseshoe loops under Nz, whereas a rounded nucleoplasm with a rounded rDNA on one side was the major morphology in Nz plus IAA (Figures 6D and 6E).

Secondly, we constructed a Cdc28-aid strain. Cells carrying *cdc28* temperature sensitive alleles, or an analog-sensitive kinase allele, arrest in G1 and become highly enlarged over time due to continuous growth ^73–75^. We depleted Cdc28 through adding IAA to the medium (Figure S6C and S6D), and found similar phenotypes (Figure 6F and S6F). In our hands, though, the *cdc28-aid* allele showed a hypomorphic phenotype when grown on YPD alone, as observed for the pseudohyphal-like growth (Figure 6F and S6F). No horseshoe rDNA loops were seen and rounded/crescent puffed Net1-GFP signals were the most common phenotype (Figure 6F). We also observed the presence of fragmented vacuoles occupying a large space, which agrees with previous findings for the *cdc28-1* allele at non-permissive temperature and for the G1 cyclin mutant *cln3Δ* ^76^. Since the Cln3p/Cdc28p complex controls vacuolar biogenesis, morphology and segregation, we cannot rule out that these underlie the absence of the horseshoe loop. However, it is more likely that Cdc28 drives loop formation through its control on cell cycle progression, condensin and/or the Nem1-Spo7/Pah1 complex previous to G2/M ^74,77–80^.

Altogether, we conclude in this chapter that the mid-M rDNA loop requires a high CDK and an inactive Cdc14.

## Discussion

The morphological reorganization of the rDNA in mitosis is one of the most remarkable cytological events in the yeast cell cycle. This is particularly outstanding during mitotic (mid-M) arrests ^9,10,13^. The nature of such reorganization has deserved multiple studies before, from the early roles of structural maintenance chromosome (SMC) complexes, mainly condensin and cohesin, to most recent works on polymer-polymer phase separation ^40,81,82^. Here, we show that growth and deformation of the nuclear envelope (NE) and its interplay with the vacuole plays a major part in the mid-M re-shaping of the rDNA (Figure 7 for a summary and model). We propose that our findings unify seemingly separated pathways that influence on the rDNA morphology. On the one hand, the aforementioned roles of cohesin and condensin and, on the other hand, the selective recruiting of newly synthesized phospholipids to the rDNA-associated NE in mid-M ^33,62^. From our data, we suggest the latter as the force that reshapes the rDNA until it becomes the outstanding horseshoe loop (Figure 7A). Thus, SMC complexes may rather provide the foundation to maintain the rDNA organized as an extensible bar during mid-M, while modulating its contraction when required; e.g., in anaphase, upon stress, etc. ^24^. Extension of rDNA bars would be favored from its spring-like configuration and the presence of locally compacted knotted domains (the observed beads; Figures 1D-H) ^81,83^, which could serve as reservoirs to nucleate extension on demand. Thus, the rDNA has the capability to extend for several microns, even when the number of units is relatively low (Figures 1B, 1C and S1). This would explain why condensin is still needed for the horseshoe loop, which is, nonetheless, the least longitudinally contracted state of the rDNA.

**Figure 7.**
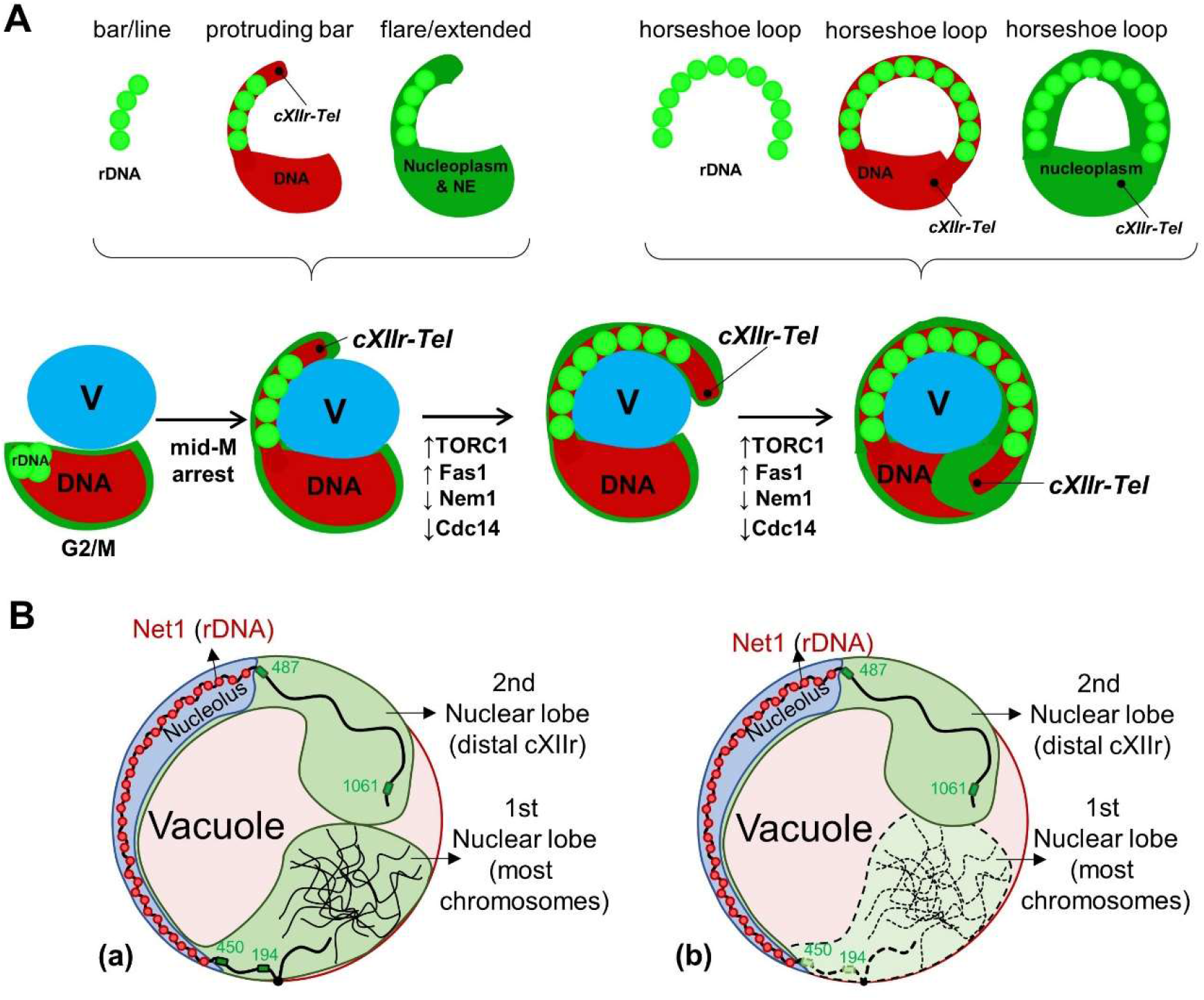
Summary and models of rDNA loops in mid-M arrests. (**A**) Relationship between rDNA morphologies (light green beads), the nuclear DNA mass– including the distal cXIIr – (red), and the shape of nucleus (dark green). On the top, the different components as seen in single and double labelling combinations, with the terminology used in this work over the schematics. The position of the cXIIr telomere is pointed. At the bottom, all components mounted together as they undergo transition towards the horseshoe loop during the Nz-driven mid-M arrest. The vacuole (V) is depicted in blue. Relative activities of key biochemical players are indicated. (**B**) The bilobed model of the horseshoe rDNA loop with the vacuole occupying the space under the loop. Both lobes are depicted separated, although they often overlap. Details of the rDNA and the whole chromosome XII are included, such as positions of the four *tetO* insertions used in this study (194, 450, 487 and 1061). Two different nucleus-vacuole spatial configurations are shown; (a) the bilobed nucleus wrap onto the vacuole; (b) the bilobed nucleus wrap around the vacuole. Dotted lines indicate structures that lay under the vacuole in (b).

Our findings also shed light on the void space that is left between the horseshoe rDNA loop and the rest of the nuclear mass (Figure 2 and S3). It is not an enlarged nucleolus as it may seem at a first glance; in fact, it is not even nuclear but occupied by one or multiple vacuoles. This means that the nucleus:cytoplasm volume ratio is not necessarily broken to attain the horseshoe loop. The intermediate states we have observed indicate that NE flares, which result from lipid biosynthesis, eventually wrap around vacuoles (Figure 3 and S4). Often, either extensive growth of these flares or mechanical pressure exerted by the vacuole makes the nucleus become bilobed, with one lobe carrying most chromosome, the second lobe just the distal part of the chromosome arm that harbors the rDNA array (cXIIr), and the handle that connects both lobes being the rDNA (Figure 7A and 7B for drawings). This spatial configuration fits well with higher order data on chromosome XII organization obtained by chromosome conformation capture (Hi-C), in which regions flaking the rDNA do not interact with each other ^84^. The horseshoe loop requires flare growth in the absence of microtubules (Nz treatment). When mid-M arrests are accomplished by means that preserve microtubules (APC inactivation or ectopic activation of SAC), other rDNA morphologies are seen (bars, lines, hooks; Figure 4 and S5). However, they can all be envisaged as incomplete horseshoe loops, obtained because spindle pulling forces pose a barrier to vacuole wrapping.

We also show that membrane phospholipid biosynthesis is required for the horseshoe rDNA loop (Figures 5G-K). It is likely that this relates to the formation of flares and expansion of the NE surface, as it has been suggested ^61–63,85^. However, phospholipid biosynthesis may be changing membrane composition of both NE and vacuoles. On the one hand, the NE appears malleable in mid-M when compared to other stages of the cell cycle and to the stiffer and spherical vacuole (Figures 3 and S4). On the other hand, the vacuolar membrane that is wrapped by the loop is poorly stained by the classical marker FM4-64 (Figure S3C). This points towards a change in the membrane composition, perhaps at specific subdomains ^86–88^, that could be crucial to establish and maintain the contacts between NE flares and the vacuolar membrane, given that known protein-protein interactions appear not to be required (Figures 6A-C). Another layer of control is defined by TORC1 itself (Figures 5A-H), which several studies relate to cell cycle progression (including signaling from mature vacuoles) and G2/M transition ^89–93^. It could be possible that TORC1 controls the localization and/or activity of specific phospholipid regulators besides the Nem1-Spo7/Pah1 complex and, likely, upregulation of lipid biosynthesis by TORC1 links the nuclear envelope (expansion) cycle with the chromosome segregation (condensation) cycle, at least in yeast ^60,87,94–97^.

The nuclear bilobed phenotype presented hereinabove resembles one atypical nuclear phenotype recently described in human cell lines, the toroidal or doughnut-shaped nucleus, in which lysosomes (vacuole equivalents in higher eukaryotes) occupy the doughnut hole ^98^. This striking morphology results from mitotic errors that stem from lysosomal impairment. Moreover, in higher eukaryotes, Lipin1 (Pah1 in yeast) is under the control of the mammalian target of rapamycin complex 1 (mTORC1) as well as the biosynthesis of lipids (FASN, ACC, etc.) through SREBP-1 ^18,99–101^. The coordination between protein and lipid synthesis is crucial for cell growth ^94,97^. Importantly, cancer cells synthesize large amounts of lipids for new membranes and, hence, for tumour growth ^102–104^. Not surprisingly, fatty acid synthase inhibitors (FASN inhibitors) are under clinical trials ^105^. Altogether, our results with yeast cells open promising new avenues for modeling these intricate processes and testing new antitumour drugs in this malleable organism.

## Materials and methods

### Yeast strains

Unless noted otherwise, all yeast strains are derivatives of W303 and YPH499 (congenic with S288C). Relevant genotypes of yeast strains used in this study are listed in Table S1. Yeast culture was performed by standard procedures. Genetic engineering was carried out through standard PCR-based procedures ^106^.

### Yeast cell growth and cell cycle

For experiments, strains were routinely grown overnight in rich YP medium (yeast extract 1% w/v + peptone 2% w/v) supplemented with 2% glucose (YPD) at 25°C with moderate shaking (150 rpm). For the standard mid-M arrest, cells were incubated with nocodazole for 180 min. First, nocodazole was added at a final concentration of 15 μg/ml, then after 120 min, half the initial concentration was added to the culture media. To arrest cells in mid-M by other means; we used either Cdc16-aid depletion or overexpression of the Mad2-Mad3 fusion protein. For Cdc16-aid depletion, cells were incubated with indole-3 acetic acid (IAA) for 180 min, at a final concentration of 5 mM. For overexpressing Mad2-Mad3, cells bearing the *PGAL1-MAD2-MAD3* construction, were grown in 2% raffinose and then overexpression from the galactose-inducible *GAL1* promoter was accomplished by growing cells in 2% raffinose + 2% galactose for 240 min. For rapamycin experiments, a final concentration of 200 nM was used (from a 2.2 mM stock in DMSO). DMSO was used at a final concentration of 1% v/v. Cerulenin was added at a final concentration of 2 μg/ml, (from a 5 mg/ml stock in EtOH). In general, indole-3 acetic acid (IAA) was added to a final concentration of 5 mM (from a 500 mM stock in DMSO), either in YPD broth or YPDA plates (YPD + agar 2% w/v).

### Pulsed Field Gel Electrophoresis (PFGE) and Southern blot

Yeast chromosomes extraction was prepared in low-melting point agarose plugs. For each sample, 6 OD_600_ equivalents were centrifuged and washed twice in ice-cold sterile 1 x PBS. Then, cells were re-suspended in Lyticase solution (2,500 Units/mL), and embedded into 0.5 % (w/v) agarose plugs. Finally, full-sized chromosomes were obtained by digesting overnight in RNaseA (10 ug/mL) and Proteinase K- (1 mg/mL) containing solutions at 37 °C. PFGE, employed to assess the chromosome XII size, was performed by using the CHEF DR-III system (Bio-Rad). 1/3 of each plug was placed within the corresponding well of a 1 % (w/v) agarose gel made in 1 x TBE buffer. Then, the wells were filled-up and sealed with additional 1 % (w/v) agarose. 0.5 x TBE was employed as the running buffer at 14 °C. The electrophoresis was carried out at 3 V/cm for 68 h, including 300 and 900 seconds of initial and switching time (respectively), and an angle of 120 °. The gel was stained with ethidium bromide for 40 minutes and destained with double-distilled water (ddH2O) for 20 minutes. The chromosomes bands were visualized under UV light using the Gel Doc system (Bio-Rad). To specifically study the chromosome XII, a Southern blot was carried out by a saline downwards transference onto a positively charged nylon membrane (Hybond-N+, Amersham-GE). A DNA probe against the NST1 region within the rDNA was synthesized using the Fluorescein-High Prime kit (Sigma-Aldrich). The fluorescein-labelled probe hybridization was carried out overnight at 68 °C. The next day, the membrane was incubated with an anti-fluorescein antibody coupled to alkaline phosphatase (Roche), and the signal was developed using CDP-star (Amersham) as the substrate. The detection was recorded by using the Vilber-Lourmat Fusion Solo S equipment.

### Western blotting

Western blotting was carried out as reported before with minor modifications ^12,107^. Briefly, 5 mL of the yeast liquid culture was collected to extract total protein using the trichloroacetic acid (TCA) method; cell pellets were fixed in 2 mL of 20% TCA. After centrifugation (2500 × g for 3 min), cells were resuspended in fresh 100 μL 20% TCA and ~200 mg of glass beads were added. After 3 min of breakage in a homogenizer (Precellys Evolution-Bertin Instruments P000062-PEVO0-A), extra 200 μL 5% TCA were added to the tubes and ~300 μL of the mix were collected in new 1.5 mL tubes. Samples were then centrifuged (2500 × g for 5 min) and pellets were resuspended in 100 μL of PAGE Laemmli Sample Buffer (Bio-Rad, 1610747) mixed with 50 μL TE 1X pH 8.0. Finally, tubes were boiled for 3 min at 95 °C and pelleted again. Total proteins were quantified with a Qubit 4 Fluorometer (Thermo Fisher Scientific, Q33227). Proteins were resolved in 7.5% SDS-PAGE gels and transferred to PVFD membranes (Pall Corporation, PVM020C-099). For protein phosphorylation states, we used the method for Phos-tag acrylamide gel electrophoresis ^108^. The following antibodies were used for immunoblotting: The HA epitope was recognised with a primary mouse monoclonal anti-HA (1:1000; Sigma); the Myc epitope was recognised with a primary mouse monoclonal anti-Myc (1:5000; Sigma-Aldrich); the Pgk1 protein was recognized with a primary mouse monoclonal anti-Pgk1 (1:5000; Thermo-Scientific) and the aid tag was recognized with a primary mouse monoclonal anti-miniaid (1:500; MBL). A polyclonal goat anti-mouse conjugated to horseradish peroxidase (1:5000, 1:10000 or 1:20000; Promega) was used as secondary antibody. Antibodies were diluted in 5% milk TBST (TBS pH 7.5 plus 0.1% Tween 20). Proteins were detected by using the ECL reagent (GE Healthcare, RPN2232) chemiluminescence method, and visualized in a Vilber-Lourmat Fusion Solo S chamber. The membrane was finally stained with Ponceau S-solution (PanReac AppliChem) for a loading reference.

### Fluorescence microscopy, staining and image processing

A Leica DMI6000B epifluorescence microscope with an ultrasensitive DFC350 digital camera was employed for single cell visualization as we have reported before ^12,109^. Briefly, 250 μl of cell culture was collected at each time point, centrifuged at 2,000 r.p.m. for 1 min at room temperature, the supernatant carefully retired, and ≈1.5 μl of the pellet was added on the microscope slide. Samples were visualized directly using a 63X/1.30 immersion objective, immersion oil with a refractive index of 1.515-1.517, and the appropriate filter cube for each tag/stain. For each field, we first captured either single planes or a series of ≈20 z-focal plane images (0.3 or 0.25 μm depth between each consecutive image), and then we processed images with the Leica AF6000 software and ImageJ. Maximum projections are indicated on figure legends. Deconvolution was performed on Z-stacks using Leica AF6000 software (method: blind deconvolution algorithm, 10 total iterations, fast processing). Orthogonal projections were generated using ImageJ. Fluorescence intensity profiles were generated using Leica AF6000 and Zeiss Zen 3.1 lite (blue edition) software. For time-lapse movies of living cells, Nz-blocked cells were pelleted and spread at a high density onto the slide. Specific conditions are described in the movie legends.

Stains: DAPI, YOYO-1, SYTO RNA Select, FM4-64, Blue CMAC. Nz-blocked cells were stained as follows:

For DAPI staining, the cell pellet was frozen for at least 24h at −20°C before thawing at room temperature, and then ≈1.5 μl of the pellet was added to ≈1 μl of 4 μg/ml of DAPI on the microscope slide.

For YOYO-1, we followed a previously described procedure ^110^. Briefly, cells were fixed in 4% formaldehyde for 30 min, washed once in PBS and re-suspended in 5 mg/ml zymolyase in P solution (1.2 M Sorbitol, 0.1 M potassium phosphate buffer pH: 6.2) for 1 min. Cells were spun down, taken up in P-Solution + 0.2% Tween 20 + 100 μg/ml RNAse A and incubated for 1 h at 37 °C. After digestion, cells were pelleted and taken up in P-Solution containing 25 μM YOYO-1, and visualized as before.

For SYTO RNASelect, cells were pelleted and stained according to manufacturer’s procedures. Briefly, a solution of RNASelect green fluorescent stain (final concentration 500 nM) in YPD medium was added to the cells and incubated for 30 min. After this, cells were washed twice with fresh YPD, let rest for 5 min and visualized as before.

FM4-64 and Blue CMAC, 1 ml of cells were grown for ≈15-30 min in either of the following: (FM4-64 final concentration of 10 ng/ml from a 5 μg/mL DMSO stock solution) or (Blue CMAC final concentration of 100 μM from a 10 mM DMSO stock solution), pelleted and visualized accordingly.

### Total Internal Reflected Fluorescence Microscopy (TIRFM)

This was adapted from a protocol described before ^111^. Nz-blocked cells were pelleted and imaged with an inverted microscope Zeiss 200 M (Zeiss, Germany) through a 1.45-numerical aperture objective (alpha Fluar, 100X/1.45, Zeiss). The objective was coupled to the coverslip using an immersion fluid (n(488) 1.518, Zeiss). The expanded beam of an argon ion laser (Lasos, Lasertechnik GmbH, Germany) was band-pass filtered (488/10 nm) and used to selectively excite EGFP-tagged proteins, for evanescent field illumination. The laser beam was focused at an off-axis position in the back focal plane of the objective. Light, after entering the coverslip, underwent total internal reflection as it struck the interface between the glass and the cell at a glancing angle. The images were projected onto a back-illuminated CCD camera (AxioCam MRm, Zeiss) through a dichroic (500 LP) and specific band-pass filter (525/50 nm). Each cell was imaged using Axiovision (version 4.9; Zeiss) with 0.5-s exposition. Image analysis: The raw images were low-pass filtered (3×3 pixels) and analysed with ImageJ.

### Confocal Superresolution Microscopy (CSM)

Nz-blocked cells were pelleted and imaged (cover glasses, high performance, D=0.17mm) in a Zeiss LSM 880 Airyscan (Axio Observer.Z1/7) super-resolution confocal inverted microscope with live cell capabilities. The resolution provided in Airyscan mode is lateral (x/y) resolution to 120 nm for 2D and 3D data sets (z-stacks) and 350-nm axial (z) resolution for z-stacks, with an improved resolution up to 1.7X compare to standard confocal ^112^. The superresolution images were taken with a C-Apochromat 63x/NA 1.20 W (Immersol Water 2010 ne=1.3339) Korr UV VIS IR M27 DICII objective. For Airyscan, the pinhole was automatically set to correct opening according to selected Airyscan mode. We used the laser line 488 (488 nm excitation wavelength and 509 nm emission wavelength) for imaging the Net1-GFP in live cells. The detection wavelength was 450-600 nm. The conditions were: Airyscan detector, power 850 V (detector gain), bidirectional scanning, channel EGFP-T3, Airyscan mode “Superresolution” (SR). For Hta2-mCherry, we used the laser line 561 (587 nm excitation wavelength and 610 nm emission wavelength). The detection wavelength was 555-700 nm. Conditions: Airyscan detector, power 728 V (detector gain), bidirectional scanning, channel mCher-T4, Airyscan mode “Superresolution” (SR). The bright field (BF) image was acquired with an ESID-T1 (transmitted light) photodiode detector, pinhole 1,00 AU. After imaging, Airyscan processing was conducted. Z-stacks were generated by applying the processing “extended depth of focus” using Zeiss Zen 3.1 (blue edition) software.

Time-lapse images on Movie 1, were acquired continuously for Net1-GFP (60 frames-1056 s). The laser line 488 (488 nm excitation wavelength and 509 nm emission wavelength) and the detection wavelength was 450-650 nm. The conditions were: Airyscan detector, power 850 V (detector gain), pinhole 5,60 AU/278 μm, unidirectional scanning (pixel time 2.33 μs, line time 9.89 ms, frame time 17.88 s), channel EGFP, Airyscan mode “Superresolution” (SR). After imaging, Airyscan processing was conducted. Specific conditions are described in the movie legends.

### Transmission Electron Microscopy (TEM)

The protocol was adapted and modified from ^113^. Cells were arrested in nocodazole for 180 min, then pelleted, re-suspended and fixed in phosphate-magnesium buffered (40 mM K_2_HPO_4_, 0.5 mM MgCl_2_ pH 6.5) 2% glutaraldehyde (EM Grade) + 2% formaldehyde and incubated overnight and stored at 4°C. Then, cells were rinsed twice in 0.1 M phosphate-citrate buffer (170 mM KH_2_PO_4_ and 30 mM sodium citrate pH 5.8) and re-suspended in this buffer containing a 1/10 dilution of Lyticase (10mg/mL 2000U stock) + Zymolyase (5U/μl stock) and incubated at 30°C for 2 h, or until cell walls have been removed. For post fixation, cells were washed twice in 0.1 M sodium acetate (pH 6.1), transferred to a 2% osmium tetroxide fixation solution and incubate for 4 hours in a fume hood. Then, cells were washed with ddH2O and transfer to 1% aqueous uranyl acetate for 60 min of incubation in the dark. After this, cells were washed twice in ddH2O and dehydrate by transferring them through a series of ethanol concentrations [20, 40, 60, 70 (overnight), 96 and 100]. Then, cells were pelleted and resuspended in EMBed 812 resin. Finally, semi-thin and ultra-thin sections were cut on an ultramicrotome (Reichert Ultracut S-Leica) and stained with toluidine blue for semi-thin sections and with uranyl acetate and lead salts [Sato’s Staining Procedure, 5 minutes] for ultra-thin sections. EM images were captured by a TEM 100 kV JEOL JEM 1010 electron microscope.

### Statistical analysis

Statistical details of experiments can be found in the figure legends.

All experiments presented in this study are representative examples, and where stated, three (n=3) or two (n=2) independent experiments (biological replicates) are shown. Number of cells counted for each condition and experiment are specified in the corresponding figure legends. Generally, more than 100 or 200 cells were counted per condition in each experiment.

For morphological data, cells were categorized and the corresponding proportions calculated and represented in bar charts. Where indicated, error bars represent Standard Error of the Mean (mean ± S.E.M).

Quantification of rDNA length and distances was performed with the Leica AF6000 software. These continuous data were represented in box-plots. In these plots, the centre lines depict the medians, box limits indicate the 25th and 75th percentile, and roughly give the 95% confidence intervals for each median. Whiskers represent 5-95 percentile. The mean is shown as ‘+’. Dots represent outliers. Exact p-values are represented on each boxplot when comparing two sets of data. For this, assumption of normality was calculated by applying a Shapiro Test. For equality of variances, an F test was used prior to a t test analysis when needed. Unpaired t test with Welch’s correction was employed for the equality of two means. When necessary, a nonparametric Wilcoxon Mann-Whitney U test was applied. Significance level was stablished at p < 0.05 (two tailed). Nonsignificant is denoted by “n.s.”

Line profile (fluorescence intensity) obtained from Leica AF6000 software in Figure S3E, was represented using GraphPad Prism.

Graph in Figure 1E represents number of beads from deconvolved images, where the central horizontal solid line represents the mean and outer horizontal solid lines represent one standard deviation (mean ± S.D.).

Quantification of bands on western blot from Figure S5H was performed by measuring the intensity of each band in non-saturated conditions using the Bio1D software. The relative amount of aid*-9myc tagged protein (target protein) was estimated using Pgk1 as an internal housekeeping control (loading control protein). Normalization of the target protein relative to the loading control protein was carried out for each lane, and then the fold difference calculated (relative target protein levels) for each lane:

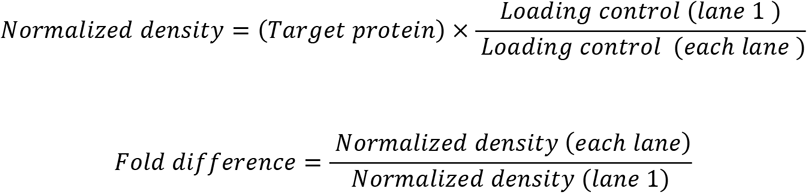

The statistical analysis was performed with GraphPad Prism (https://www.graphpad.com/) and R software (https://www.r-project.org/).

## Acknowledgments

We kindly thank the Frank Uhlmann lab and the Mike Henne lab for yeast strains. We thank the José Manuel Siverio lab for reagents. We thank María Moriel-Carretero for personal communication of unpublished results. We also thank David Machado from the Pharmacology Unit at Universidad de La Laguna for support on TIRF microscopy and José Manuel Pérez Galván from Servicio Investigación Microscopía Avanzada Confocal y Electrónica – SIMACE – at Universidad de Las Palmas de Gran Canaria for support on confocal and electron microscopy.

This work was supported by the Spanish Ministry of Science grant BFU2017-83954-R to F.M.

The authors declare no competing financial interests.

## Author contributions

Conceptualization, E.M-P., and F.M.; Methodology, E.M-P., S.S-S., and F.M., (all strains stated as “this work” in table S1 created by E.M-P, except for strain FM2743, created by S.S-S.); Formal Analysis, E.M-P., J.A-P., S.S-S., and F.M.; Investigation, E.M-P. (all experiments unless noted otherwise), J.A-P. (experiments in Figures 1C and S1), S.S-S., and F.M. (experiments in Figures 3A and Movies 2-4), F.M. (experiment in Figure S3A); Writing – Original Draft, E.M-P., and F.M.; Writing – Review & Editing, E.M-P., and F.M.; Visualization, E.M-P., J.A-P., and F.M.; Supervision, F.M.; Project Administration, F.M.; Funding Acquisition, F.M.

## MOVIE LEGENDS

**Movie 1.** The rDNA loop is static in its length and curvature while highly dynamic across. A Net1-GFP horseshoe-like loop was selected from a mid-M arrest (strain FM931) and filmed under a superresolution mode with an LSM800 Airyscan confocal microscope (60 cycles of 18 sec). Cells were quickly spread on a slide, and filmed immediately after placing the coverslip. In these conditions, the loop length remains static for the first 10 min (~600 sec). After that, longitudinal contraction is observed. Two movie frames (125 and 233 secs) are shown and further commented in Figure 1G and H.

**Movie 2.** The chromosome XII right arm (cXIIr) telomere is highly mobile in the mid-M arrest. The strain carrying the *tetO:450* (rDNA proximal flank) *tetO:1061* (cXIIr-Tel) TetR-YFP Net1-eCFP (FM2301) was filmed as in movie 1 but with the epifluorescence microscope DMI6000 (59 cycles of 10 sec). A montage of the TetR-YFP channel, the Net1-eCFP channel (pseudo-coloured in red) and a merge of both is shown. Note that the *tetO:450* remains static and attached to one rDNA edge while the *tetO:1061* rapidly moves around. We know which is which because a strain carrying *tetO:450* and *tetO:487* (distal rDNA flank) Net1-eCFP (FM2438) statically maintains both *tetOs* at each Net1-eCFP flank.

**Movie 3.** The dynamics of morphologically diverse rDNA loops (I). The strain carrying Net1-GFP Hta2-mCherry was filmed as in movie 2, although with frames taken every 30 sec instead of 10 sec. A montage of the BF, Net1-GFP and Hta2-mCherry channels as well as the merge of Net1 and Hta2 is shown. The cell confluency on the slide was very high in order to film as many nuclei as possible, at the expense of information on individual cells (see BF; however, >95% were mononucleated dumbbell shaped cells, as expected for a mid-M arrest). Fifteen nuclei are numbered. Most of them carry horseshoe rDNA loops, yet seen under different visual angles: front, or almost front, views (#1,2,4,6,10,13), top views (#8, 15) and side views (#5). Nucleus #3 changed from a front to a top view through the time course. Other rDNA morphologies are more difficult to assign. For example, the nucleus #7 (and even #5 in some frames) appears as a protruding bar; whereas nuclei #9,12 and 14 may correspond to ovals seen under different angles. From 5-10 min onwards, loops are longitudinally contracted almost synchronously. Contraction is across the loop rather than by flare/bar recoiling from one edge, so that the horseshoe appearance is maintained and the SUL is reduced. Note that flanking rDNA sequences dynamically become part of the loop (e.g., #2,4,10 and 13; the latter indicated by an arrowhead). Note also that Hta2-mCherry appears bipartite in top-view loops (#8 and #15, the latter indicated by an asterisk), which agrees with the bi-lobed nucleus model.

**Movie 4.** The dynamics of morphologically diverse rDNA loops (II). Like in movie 3, but with frames every 10 sec. Five nuclei are numbered, all carrying horseshoe rDNA loops seen under different visual angles: front (#1,2) and top views (#3,4,5). As in movie 1, loops are static in length and curvature during the first 10 min.

