## Supplemental Figures and Tables for "Nuclear envelope reshaping around the vacuole determines the morphology of the ribosomal DNA"

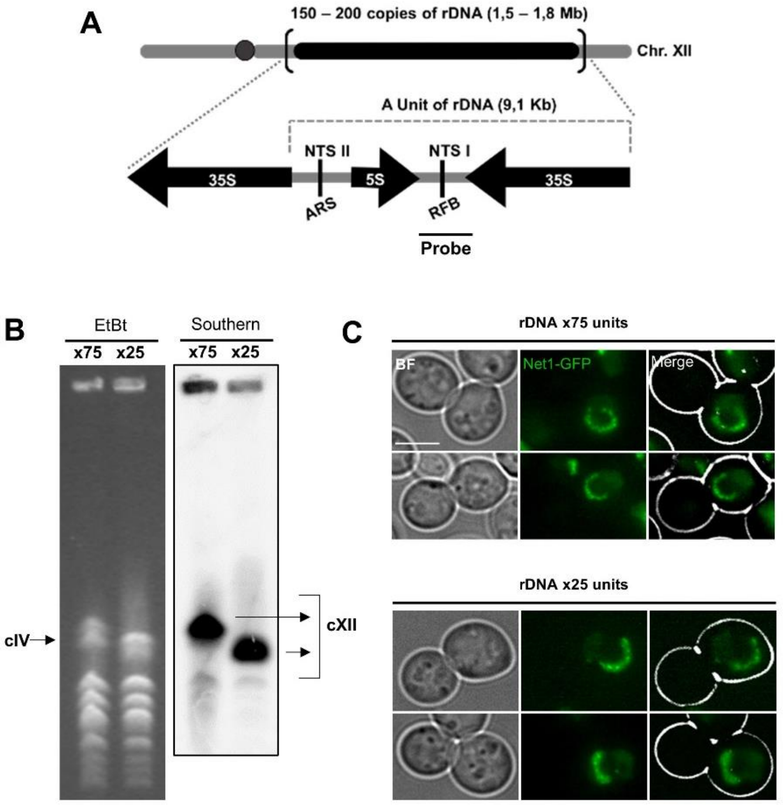


**Figure S1. Confirmation of distinct rDNA lengths used to check their influence in the size of the rDNA loop.** Related to Figures 1B and 1C.

(**A**) Unscaled drawing of yeast chromosome XII, with the canonical 150-200 copies rDNA array indicated by a black bar on the right arm. Genetic details of the single repetitive unit are shown below, together with the position and length of the probe used in the Southern blots. 35S and 5S, genes for the corresponding pre-rRNAs; NTS I & II, non-transcribed sequences I & II; ARS, autonomous replication sequence; RFB, replication fork block.

(**B**) Confirmatory pulsed-field gel electrophoresis (PFGE) of two clones with ~75 and ~25 copies rDNA arrays. Running conditions are optimal for maximizing separation by size of chromosome XII (cXII) with rDNA arrays ranging from 2 to >200 copies [1]. On the left, the ethidium bromide staining of the pulsed-field gel. The band corresponding to chromosome IV (cIV), the largest yeast chromosome with a fixed size (1.53 Mbps), is indicated. The cXII is actually larger than cIV when the rDNA array is greater than 50 copies. The x25 lane correspond to a Δ*fob1* strain previously constructed with 25 copies [2]. Accordingly, the size of cXII is smaller than cIV and the cXII band migrates faster than that of cIV. The x75 lane correspond to a Δ*fob1* clone that originates from a x190 strain that spontaneously suffered from a Fob1-independent shortening of the array. Accordingly, its cXII band migrates slightly slower than cIV. On the right, Southern blot of the PFG with a probe against the rDNA to confirm the cXII bands.

(**C**) Examples of mid-M rDNA loops from both strains. Quantification of loop lengths are in Figure 1C. Scale bar is 5 µm.

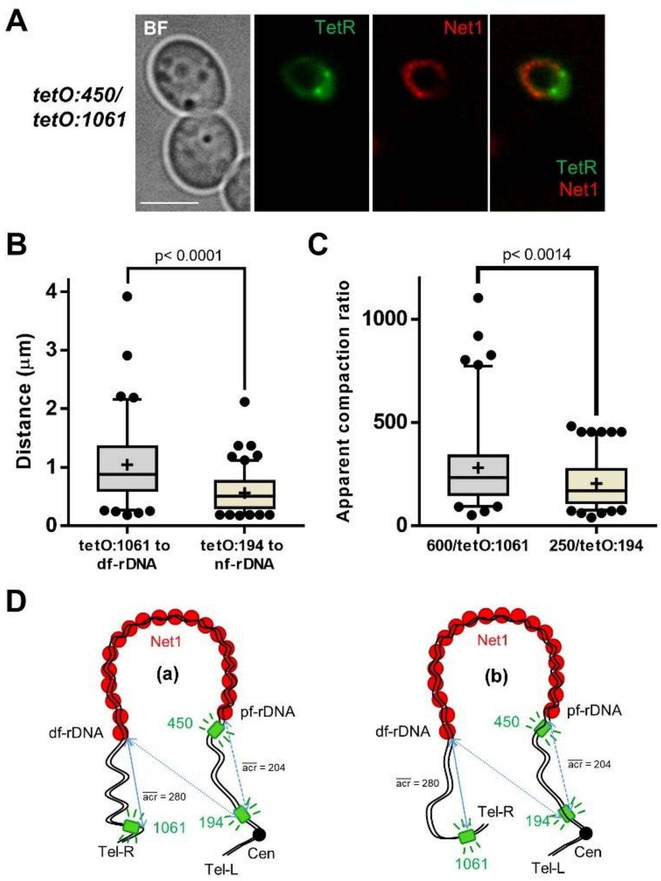


**Figure S2. Putative configurations of the chromosome XII in the rDNA loop.** Related to Figures 1I and 1J.

(**A**) The horseshoe rDNA loop in a strain with both 450 (proximal rDNA flank) and 1061 (right arm telomere) *tetOs*, the TetR-YFP and the Net1-eCFP (pseudo-colored in red). Note that one *tetO* (likely the 450) is close and attached to one base of the loop, whereas the second *tetO* (likely the 1061) is close, yet not attached, to the second base. The scale bar represents 5 µm.

(**B**) Quantifications of the distance between the *tetO:1061* and the distal flank of the rDNA (df-rDNA; identified as the one that did not have the *tetO:450* attached) (n=102 cells), as well as the distance between the *tetO:194* (centromere) and its nearest rDNA flank (nf-rDNA; it is not possible to assess whether the nf-rDNA corresponds to the pf-rDNA or the df-rDNA) (n=120 cells). Boxplot: Whiskers represent 5-95 percentile. Mean shown as '+'. Dots represent outliers.

(**C**) Conversion of the data shown in (B) into apparent compaction ratios for the centromere to pf-rDNA and df-rDNA to right arm telomere tracts. Boxplot: as in B.

(**D**) Schematics of non-mutually exclusive spatial configurations for the chromosome XII in the horseshoe rDNA loop, as deduced from (C): (a), the rDNA-telomere tract is curlier than the centromere-rDNA tract, giving the impression of an apparent higher compaction ratio; (b) the rDNA-telomere tract folds back, with similar effects on the apparent compaction. The centromere-rDNA tract might adopt a configuration that tends to maximize distance (acr = 204; not far from 140, the reported mitotic compaction ratio [3]). See later figures in this manuscript for further refinement on these configurations. The green boxes indicate the *tetO* arrays at 194 (centromere), 450 (rDNA proximal flank; pf-rDNA) and 1061 (right arm telomere). The solid blue lines indicate the distance between the telomere and the df-rDNA. The dotted blue lines indicate the two possible distances measured for the centromere without a reference for either rDNA flank.

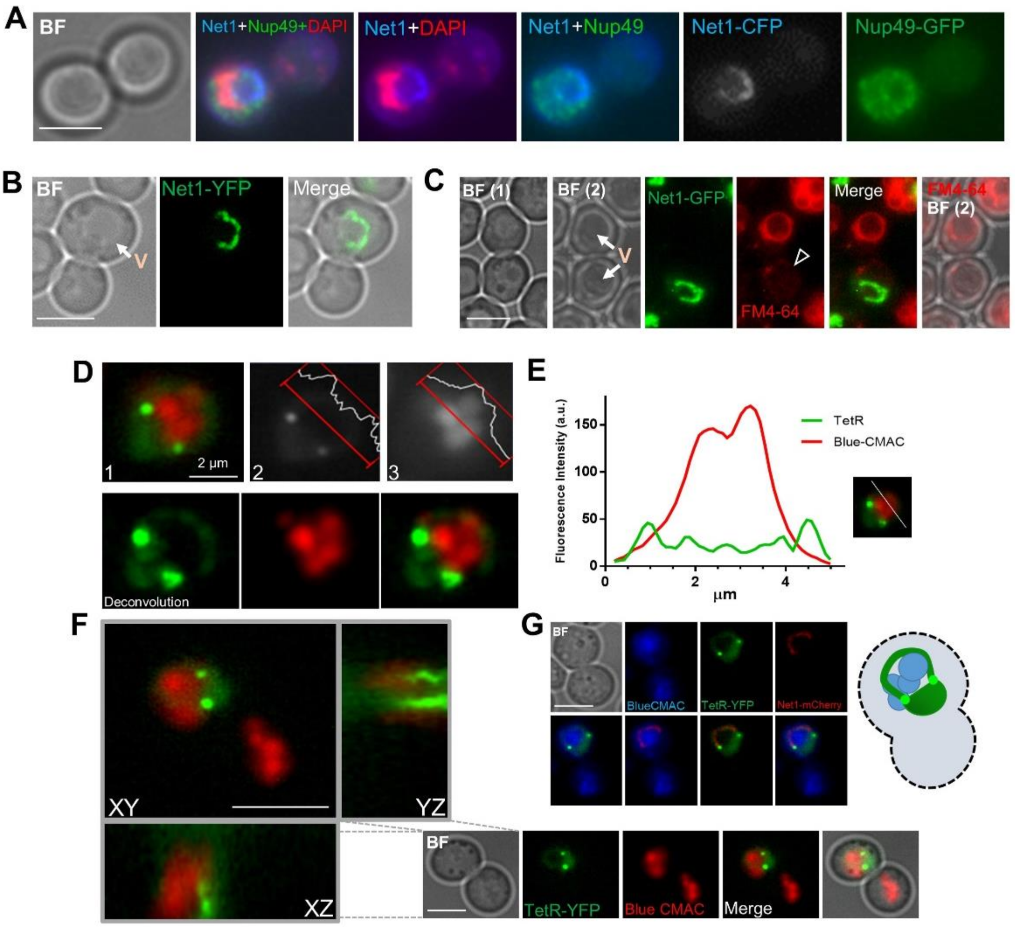


**Figure S3. The space under the rDNA loop (SUL) is not nuclear and contains vacuole(s).** Related to Figure 2.

(**A**) The nuclear envelope marker Nup49 weakly stains the SUL in horseshoe rDNA (Net1) loops. Unlike Figure 2I, these samples were processed for DAPI staining of the nuclear mass.

(**B**) Example of an rDNA loop that crosses a vacuole (V; pointed by an arrow in the BF). This is likely due to the spatial orientation of both (the loop is partly seen from the top). Note that the loop edges touch the vacuole circumference. See Figure 2L for a horseshoe loop that almost perfectly follows the vacuole curvature.

(**C**) The vacuolar membrane (VM) marker FM4-64 does not stain the VM of vacuoles associated with rDNA loops. Note how vacuoles of similar size (V; pointed by white arrows) are distinctly stained by FM4-64; the one with the rDNA loop is stained very weakly (hollow arrowhead). To better appreciate both Vs, top and bottom out of focus BF images are shown.

(**D-G**) The SUL is fully occupied by vacuolar content. (**D**) An example of the bag-like nucleus that leaves a horseshoe loop with the vacuole lumen occupying the SUL. The strain bears tetO:450 tetO:1061 TetR-YFP, and the vacuole lumen is stained with Blue CMAC (pseudo-colored in red in the RGB pictures). (1) Maximum projection of the montage of channels; (2) the TetR-YFP channel (grayscale); (3) the Blue CMAC channel (grayscale). Intensity profiles of a SUL cross section are shown for each channel. (**E**) Chart of the intensities across the SUL. Note how two peaks of nucleoplasm intensity (in green) lay at the borders of the vacuolar lumen (in red). (**F**) Orthogonal projections of a second example of a horseshoe loop with the vacuole in the SUL. Note how *tetOs* lay on different Z planes. This is rather common in front views of horseshoe loops. (**G**) A third example of a horseshoe loop with the vacuole in the SUL. In this case, the mid-M cell carries another major vacuole in the second cell body. The nucleus only interacts with the vacuole of its same cell body.

Scale bars represent 5 µm unless stated otherwise. BF, bright field.

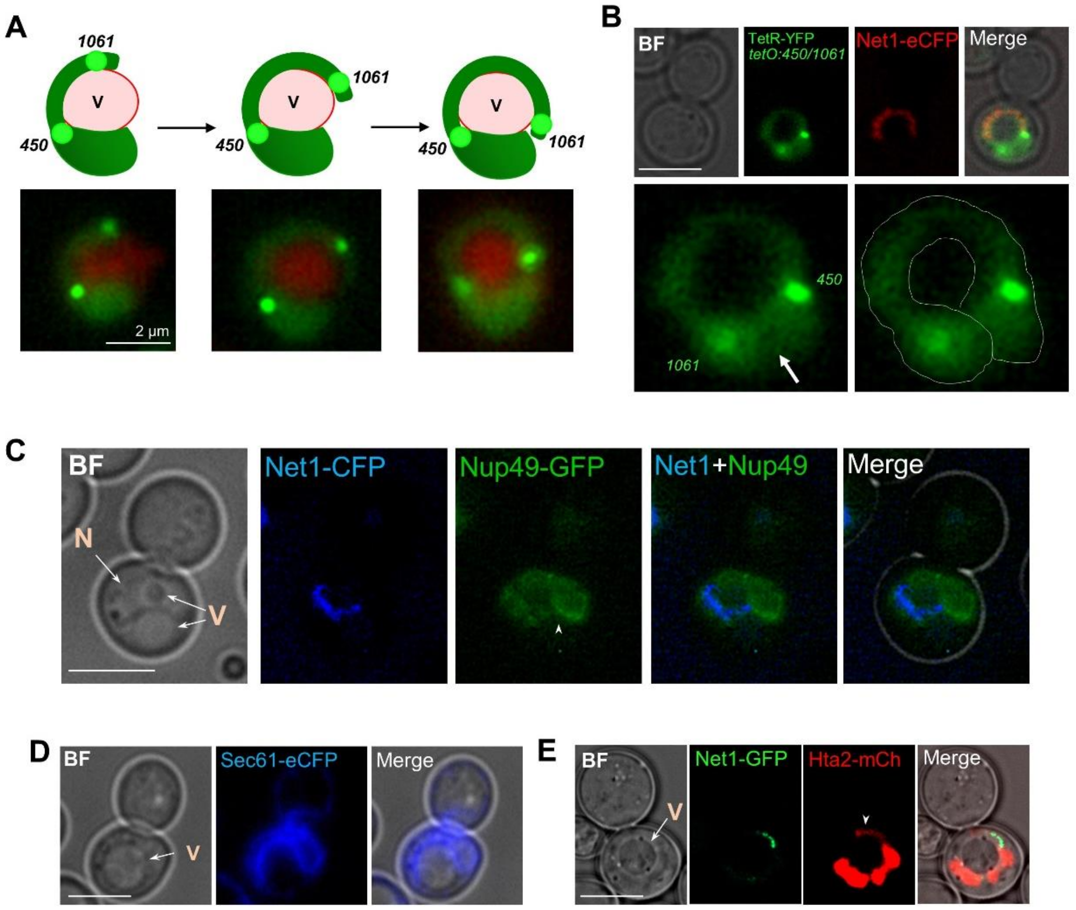


**Figure S4. The horseshoe rDNA loop is formed when the nuclear flare that contains the protruding rDNA bar folds back and its tip overlaps with the main nucleus.** Related to Figure 3.

(**A**) The rDNA bars within the nucleoplasm flares can be seen as precursor of the horseshoe loop. The drawings on the top represent nuclear flares as they originate and bend around the vacuole. The nucleoplasm is in dark green, the *tetOs* spots are in light green, and the vacuole in light orange (lumen) and red (vacuolar membrane). Under each schematics of growing flares, an actual example taken from the strain with the *tetO:450 tetO:1061* TetR-YFP Net1-eCFP in which the vacuolar lumen has been stained with Blue CMAC (pseudo-colored in red).

(**B**) The horseshoe rDNA loop often comprises a bilobed nucleus, with the rDNA being the handle. Another example taken from (A) is shown. In this case, bending is complete and the distal cXIIr has become lobed and overlaps with the lobe that carries the rest of cXII. The *tetO:1061* is recognized because it is not closely attached to one edge of the Net1 loop, as is the *tetO:450*. Details of the bended bi-lobed nucleus are shown in the lower TetR-YFP micrographs. The arrow on the left image points to the constriction that separate both overlapping lobes. On the right, a solid white line has been drawn to indicate the spatial configuration of the overlapping lobes. The YFP and CFP channels correspond to a single z plane. In all examples of this kind of horseshoe loop where the Net1-eCFP is seen throughout (i.e., perfect frontal view), the tetO:450 is on focus and the tetO:1061 is out of focus, further supporting the bi-lobed model.

(**C**) An example taken from the Net1-CFP Nup49-GFP strain where the flare folds back and the tip touches the NE of the main nucleus (pointed by an arrowhead). Note that in the BF two vacuoles (V) can be envisioned; the flare bends around the one on the top, whereas the one at the bottom also influences on the nuclear/flare morphology through deformation. This BF also shows a rare example in which the nucleus (N) can also be envisioned (compare BF and Nup49-GFP).

(**D**) An example of a nucleus deformed by localizing between a large vacuole (V) and the cell surface. The nucleus is visualized through the nER marker Sec61-eCFP. While growing nuclear flares may explain many of the observations reported here, nuclear deformation by squeezing between stiff vacuoles and the cell surface could explain others. It might even aid flare formation and grow.

(**E**) Another example of nuclear deformation by the vacuole as seen by CSM. In this case, most of the nucleus, not only the rDNA bar (pointed by an arrowhead), bends around the vacuole (arrow). The nuclear DNA mass (Hta2-mCherry) appears unevenly distributed across the bended nucleus.

Scale bars represent 5 µm unless stated otherwise. BF, bright field.

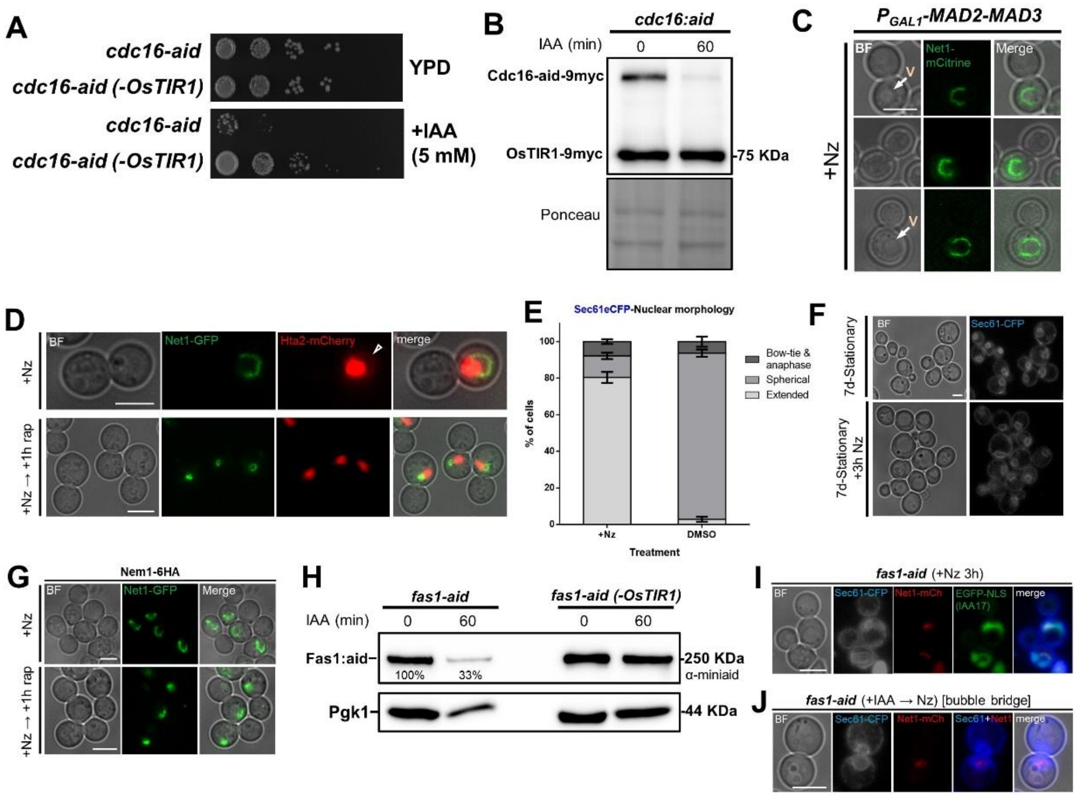


**Figure S5. Confirmation of distinct genetic modification used to check mechanisms of rDNA loop formation, and effects of TORC1 on the rDNA loop.** Related to Figures 4 and 5.

(**A**) Spot assays to check Cdc16-aid degradation under auxin. Cdc16 is an essential protein. A strain carrying a cdc16-aid allele plus the auxin-mediated degron system (OsTIR1) was spotted with and without auxin (indol acetic acid; IAA) alongside a strain derivative where OsTIR1 has been eliminated. Note that cdc16-aid OsTIR1 does not grow on YDP plus 5 mM IAA

(**B**) Western blot (WB) to check Cdc16-aid degradation upon IAA addition. Samples were taken before (0) and 60' after adding 5 mM IAA. The WB (upper picture) is against the myc epitope, which is in both Cdc16-aid (-9myc) and OsTIR1 (-9myc). The latter serves as an internal control. Total protein loading is shown in the lower picture after Ponceau staining.

(**C**) Examples of cells carrying the $P_{GAL1}$-*MAD2-MAD3* construction after being arrested with Nz for 3h in glucose. The aim of this experiment was to check that the horseshoe rDNA loop is present in this particular genetic background.

(**D**) TORC1 inactivation in mid-M turns the horseshoe loop into oval, short lines or hyper mini-loops. Representative cells upon Nz alone or Nz followed by rapamycin (1h). The arrowhead points to a Hta2 handle.

(**E**) Quantification of NE morphologies in either asynchronous cultures treated with DMSO 1% v/v for 3h or arrested with Nz for the same period (Nz treatment also implies having DMSO 1% v/v in the cultures) (mean ± S.E.M., n=3). See Figure 5D for more details.

(**F**) Nz does not prompt NE flares in stationary phase (7 days of continuous growth in a flask).

(**G**) Microscopy of the samples taken for the Western blots of Figure 5G (Nem1-HA). Note how the horseshoe loop in Nz gets contracted after TORC1 inactivation (rapamycin addition), as we have shown before [4].

(**H**) WB to check Fas1-aid degradation upon IAA addition. Samples were taken before (0) and 60’ after adding 5 mM IAA. The upper WB is against the aid epitope; the lower WB is against Pgk1 (reference). Two strains were checked, one with Fas1-aid plus OsTIR1 (left) and a derivative where OsTIR1 has been removed (right).

(**I**) Control check that C-tagging of Fas1 with aid does not interfere with loop formation upon Nz. This strain carries the nucleoplasm marker EGFP-NLS, which also contains the auxin degron IAA17 sequence, related to aid. This marker disappears after IAA treatment. Note the bilobed EGFP-NLS signal.

(**J**) Example of the NE bubble bridge category, seen as a minor outcome in Fas1-aid upon IAA. This category is closely related to the bow-tie, but with more spherical dumbbell-like appearance.

Scale bars represent 5 µm. BF, bright field; V, vacuole.

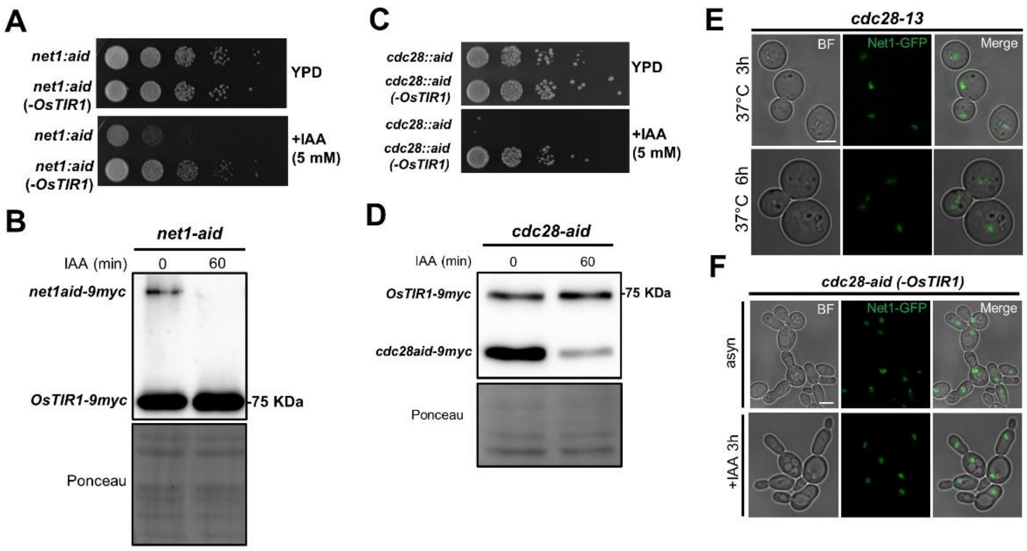


**Figure S6. Confirmation of distinct genetic modifications used to check cell cycle influence on the rDNA loop.** Related to Figures 6.

(**A**) Spot assays to check Net1-aid degradation under auxin. The procedure is described in Figure S5A.

(**B**) WB to check Net1-aid degradation upon IAA addition. The procedure is described in Figure S5B.

(**C**) Spot assays to check Cdc28-aid degradation under auxin. The procedure is described in Figure S5A.

(**D**) WB to check Cdc28-aid degradation upon IAA addition. The procedure is described in Figure S5B.

(**E**) Cell and nucleolar morphology (Net1) in *cdc28-13* (thermosensitive allele for *CDC28*) at restrictive conditions (37 ºC). The purpose of this experiment is to compare cell morphology with *cdc28-aid* under restrictive conditions (Figure 6F). Note that cells become enlarged in both cases. The *cdc28-13* was not used in this study because of the effects of the HS on the rDNA.

(**F**) C-terminal tagging of Cdc28 with aid makes cells grow as chains. This complements the results from Figure 6F by showing that this phenotype is due to the *cdc28-aid* allele alone (not the combination of *cdc28-aid* and OsTIR1).

Scale bars represent 5 µm. BF, bright field.

### Table S1. Strains used in this work.

| Figure | Strain[1] | Genotype[2] | Origin |
|---|---|---|---|
| – | AS499 | *MATα ura3-52 lys2-801 ade2-101 trp1-Δ63 his3-Δ200 leu2-Δ1 bar1-Δ* | Strunnikov lab |
| **1D** | CCG743 | {AS499} *SIR2-GFP::LEU2* | Aragon lab |
| **1A** | CCG771 | {AS499} *NET1-GFP::LEU2* | Aragon lab |
| **1D** | CCG918 | {AS499} *CDC14-GFP::KanMX* | Aragon lab |
| – | CCG1297 | {AS499} *TetR-YFP::ADE2 TetO(5.6Kb)::194Kb-ChrXII::HIS3* | Aragon lab |
| – | CCG1300 | {AS499} *TetR-YFP ADE2 TetO(5.6Kb)::487Kb-ChrXII::HIS3* | Aragon lab |
| **2I; S3A; S4C** | CCG1582 | {AS499} *NET1-CFP::HygB; NUP49-GFP::URA3* | Aragon lab |
| – | CCG2306 | {AS499} *TetR-YFP::ADE2 TetO(5.6Kb)::450Kb-ChrXII::URA3 TetO(5.6Kb)::487Kb-ChrXII::HIS3* | Aragon lab |
| **2M; 3H-I; S3D-F; S4A** | CCG2309 | {AS499} *TetR-YFP::ADE2 TetO(5.6Kb)::450Kb-ChrXII::URA3 TetO(5.6Kb)::1061Kb-ChrXII::HIS3* | Aragon lab |
| **1C; S1** | CCG2570 (x75) | *MATa leu2-3,112 ura3-1 his3-11 trp1-1 ade2-1 can1-100 fob1Δ::his3::HygB (rDNA ~ 75 copies); NET1-GFP::LEU2* | Aragon lab |
| **1C; S1** | CCG2572 | *MATa leu2-3,112 ura3-1 his3-11 trp1-1 ade2-1 can1-100 fob1Δ::his3::HygB (rDNA ~ 25 copies); NET1-GFP::LEU2* | Aragon lab |
| – | SEY6210 | *MATα ura3-52 leu2-3,112 his3-Δ100 trp1-Δ901 lys2-801 suc2-Δ9* | Henne lab |
| – | SEY6210 mdm1Δ nvj3Δ | {SEY6210} *mdm1Δ::KanMX nvj3Δ::NatMX (MATα)* | Henne lab |
| – | SEY6210 ΔNVJ | {SEY6210} *nvj1Δ::TRP1 nvj2Δ::HIS3 mdm1Δ::KanMX nvj3Δ::NatMX (MATα)* | Henne lab |
| **4B,E,F,H; S5C** | W303-K699 | *MATa trp1-1 can1-100 leu2-3,112, his3-11,15, ura3-1 GAL phi+ ade2-1::OsTir1-9Myc::ADE2 smc4-3HA::TRP1 Net1-yEmCitrine::HIS3 leu2-3,112::*$P_{GAL}$*-MAD2-MAD3::LEU2* | Uhlmann lab |
| – | YAT1735 | *MATα cdc28-13 leu2 his ura3 trp1* | Toh-e lab[3] |
| – | yED233 | *Mata ura3-1 HTA2-mCherry::URA3 ade2-1 his3-11,15 leu2-3,112 trp1-1 can1-100* | Pelet lab |
| **2C,E** | YNK54 | *Mata ura3-1::ADH1-OsTIR1-9Myc::URA3 ade2-1 his3-11,15 leu2-3,112 trp1-1 can1-100* | Kanemaki lab |
| **1D-H; 2J; 3G; 5E; S3B,C** | FM931 | {YNK54} *NET1-GFP::LEU2* | Machín lab |
| **2B** | FM2113 | {YNK54} *SPC42-RedStar::KanMX [NOP1-CFP(LEU2)]* | Machín lab |
| **2G; 3E; S2A-C; S4B** | FM2301 | {CCG2309}; *NET1-eCFP::KanMX4* | This work |
| **2A; S2B,C** | FM2361 | {CCG1297}; *NET1-eCFP:KanMX4* | This work |
| **5G,H; S5G** | FM2383 | {FM931}; *NEM1-6HA::natNT2* | This work |
| **5D,F; S4D; S5E,F** | FM2394 | {YNK54}; *SEC61-eCFP:kanMX4* | This work |
| **4B-D,G; S5A,B** | FM2396 | {FM931}; *cdc16-aid*-9myc::hphNT* | This work |

| | | | |
|---|---|---|---|
| **6D,E** | FM2398 | {CCG1300} *ura3-52::ADH1-OsTIR1-9Myc::URA3; net1-aid*-9myc::hphNT; FOB1-eCFP::KanMX4* | This work |
| **1J** | FM2438 | {CCG2306}; *NET1-eCFP:KanMX4* | This work |
| **2D,L; 3A-D; 5A-C; S4E; S5D** | FM2614 | {yED233}; *NET1-GFP::LEU2* | This work |
| **S6E** | FM2619 | {YAT1735}*; NET1-GFP::LEU2* | This work |
| **S6A** | FM2620 | {FM2398} *ura3-52 (-OsTIR1)* | This work[4] |
| **2H; 3K** | FM2639 | {FM2394}; *NET1-mCherry::natNT2; trp1-1::$P_{ADH1}$-EGFP-IAA17-NLS::TRP1* | This work[5,6] |
| **2K** | FM2641 | {FM2394}; *NET1-mCherry::natNT2; trp1-1::$P_{ADH1}$-EGFP-IAA17::TRP1* | This work[5,6] |
| **3J; S3G** | FM2657 | {CCG2309}; *SEC61-eCFP:kanMX4; NET1-mCherry:natNT2* | This work |
| **S5A** | FM2659 | {FM2396}; *ura3-1 (-OsTIR1)* | This work[4] |
| **6F; S6C,D** | FM2694 | {FM931}; *cdc28-aid*-9myc::hphNT* | This work |
| **S6C,F** | FM2695 | {FM2694} *ura3-1 (-OsTIR1)* | This work[4] |
| **6A-C** | FM2696 | {SEY6210}; *NET1-GFP::LEU2* | This work |
| **6A-C** | FM2697 | {SEY6210 mdm1Δ nvj3Δ}; *NET1-GFP::LEU2* | This work |
| **6A-C** | FM2698 | {SEY6210 ΔNVJ *(nvj1Δ,nvj2Δ,mdm1Δ,nvj3Δ)*}; *NET1-GFP::LEU2* | This work |
| **2F; 3F** | FM2743 | {yED233}; *Sec61-EYFP::kanMX4* | This work |
| **5I-K; S5H-J** | FM2800 | {FM2639}*; fas1-aid*-9myc::hphNT* | This work |
| **S5H** | FM2801 | {FM2800} *ura3-1 (-OsTIR1)* | This work[4] |

[1] Strains are sorted alphabetically, and then by number, starting from strains reported in previous works.

[2] Curly brackets indicate parental strains used for successive strain construction. Semicolons separate independent transformation events during strain construction; intermediate strains are omitted. Square brackets indicate episomal elements.

[3] These strains were obtained from the NBRP repository (http://yeast.lab.nig.ac.jp/yeast/).

[4] These strains were obtained by counterselecting for the ura$^{-}$ phenotype in 5-FOA, which results in the pop out of the OsTIR1::URA3 segment.

[5] These strains express EGFP reporters for the nucleoplasm and the cytoplasm, respectively. These EGFPs are chimeras that contain the auxin-responsive degron peptide IAA17 from *Arabidopsis thaliana*. In most figures, the reference to the IAA17 is omitted for the sake of space since it is experimentally irrelevant. In those cases where cells were treated with IAA, the IAA17 epitope is indicated.

[6] The integrative plasmids for making these strains were also obtained from the NBRP repository (pMK42 and pMK72; originally from Kanemaki lab). To target pop-in integration into the *TRP1* locus, both plasmids were digested with MfeI prior to transformation.

**Supplemental references.**


1. Machín, F., Torres-Rosell, J., De Piccoli, G., Carballo, J.A., Cha, R.S., Jarmuz, A., and Aragón, L. (2006). Transcription of ribosomal genes can cause nondisjunction. J. Cell Biol. *173,* 893–903.

2. Kobayashi, T., Heck, D.J., Nomura, M., and Horiuchi, T. (1998). Expansion and contraction of ribosomal DNA repeats in Saccharomyces cerevisiae: requirement of replication fork blocking (Fob1) protein and the role of RNA polymerase I. Genes Dev. *12,* 3821–3830.

3. Guacci, V., Hogan, E., and Koshland, D. (1994). Chromosome condensation and sister chromatid pairing in budding yeast. J. Cell Biol. *125,* 517–30.

4. Matos-Perdomo, E., and Machín, F. (2018). The ribosomal DNA metaphase loop of Saccharomyces cerevisiae gets condensed upon heat stress in a Cdc14-independent TORC1-dependent manner. Cell Cycle *17,* 200–215.
